# A logratio approach to the analysis of autosomal genotype frequencies across multiple samples

**DOI:** 10.1101/2025.01.17.633675

**Authors:** Jan Graffelman

## Abstract

John Aitchison has shown that the logratio principal component analysis of multiple samples of a biallelic polymorphism can evidentiate the Hardy-Weinberg law. This article extends Aitchison’s work to multiallelic polymorphisms showing how the Hardy-Weinberg law manifests itself in a logratio based statistical analysis of larger genotypic compositions. Some fundamental relationships between allelic and genotypic compositions are derived, and the close relationships between the logratio principal component analysis of allelic and genotypic compositions and the corresponding compositional biplots are established. We perform simulations and practical genetic data analysis to explore the implications of Hardy-Weinberg equilibrium for the logratio principal component analysis of genotypic compositions. A general multiallelic compositional measure for disequilibrium is presented, and shown to relate to the classical inbreeding coefficient. The proposed compositional analysis is illustrated with biallelic glyoxalase genotypes and multiallelic forensic STRs from the 1,000 Genomes project.

## 1 Introduction

The Hardy-Weinberg equilibrium (HWE) is a well-known foundational principle of population genetics (Hardy, 1908; Weinberg, 1908). For a biallelic genetic polymorphism with alleles A and B with respective frequencies *p* and *q*, the random combination of alleles implies the relative expected genotype frequencies *f*_*AA*_, *f*_*AB*_ and *f*_*BB*_ are *p*^2^, 2*pq* and *q*^2^ respectively. Squaring the heterozygote frequency, this can be rephrased as a compositional law that relates the three parts of a genotypic composition:

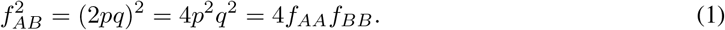

More broadly, this is a particular case of a power law given by 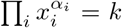 The equation is linearized by taking logarithms

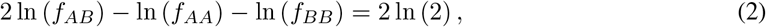

which shows the law in its *logcontrast form* (see Section 2.1). The latter two equations clearly establish a link between population genetics and the statistical field of compositional data analysis (CoDA (Aitchison, 1986)), when genotype and allele frequencies are considered as compositions. Graphical representations of three-part compositions in ternary diagrams, like in Figure 1, are widely used in CoDA and other scientific disciplines. De Finetti (1926) represented a biallelic genotypic composition in a ternary diagram; the latter is also known as a *de Finetti diagram* in the genetics literature (Cannings and Edwards, 1968; Edwards, 1977). Compositions in HWE are described by a parabola in the ternary diagram. The AA-BB base of the diagram represents an axis for the allele frequency of the composition (see Fig. 1A). The significance of deviations from the equilibrium parabola can be judged by representing the acceptance region of a statistical test for equilibrium in the ternary diagram (Graffelman and Morales-Camarena, 2008). Aitchison (1999), in one of his multiple seminal contributions to the analysis of compositional data, showed how a natural law like HWE manifests itself if a set of biallelic genotypic compositions is analysed by logratio principal component analysis (LR-PCA; Aitchison (1983)); the law can be inferred from the eigenvector coefficients obtained in LR-PCA (see Section 3.1).

**Figure 1:**
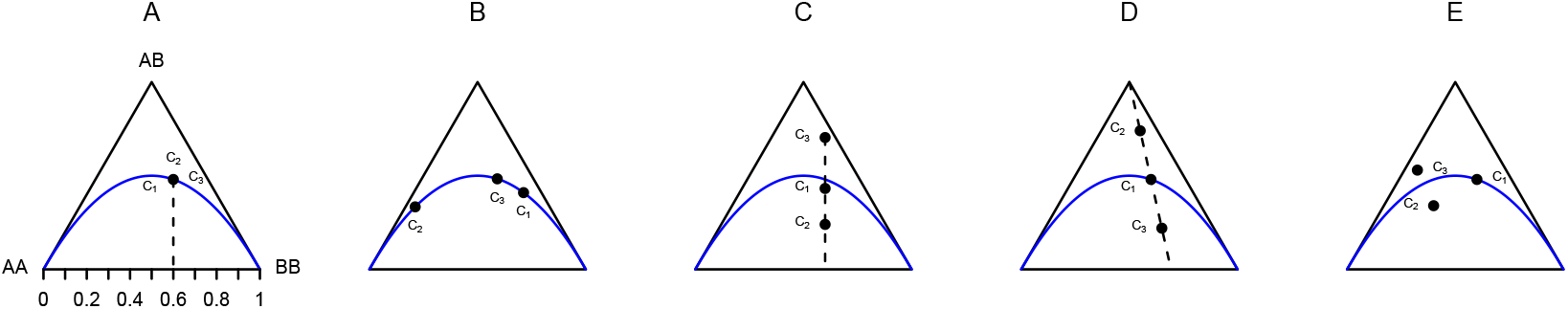
Ternary diagrams with three biallelic genotypic compositions. A: Equilibrium compositions with no variation in allele frequencies or ratio of homozygotes. B: Equilibrium compositions with variation in allele frequencies. C: Compositions with constant allele frequency and variation in the degree of disequilibrium. D: Compositions with a constant ratio of the homozygotes AA and BB. E: Compositions with variation in allele frequencies, in ratio of homozygotes and in degree of disequilibrium. In panel E, composition *C*_1_ is an equilibrium composition, *C*_2_ has a lack of heterozygotes and *C*_3_ an excess of heterozygotes.

The purpose of this article is to expand Aitchison’s work to the case of multiple alleles, and to explore how HWE manifests itself in the LR-PCA of both biallelic and multiallelic genotype compositions, by means of both simulation and practical genetic data analysis. We extend the ratio-based disequilibrium measures for HWE that have been used in the statistical genetics literature (Ziegler and König, 2006). We will consider multiple samples, all taken for the same genetic variant, as genotypic compositions, resulting from the division of the genotype counts by the sample size. Such data is obtained either by repeatedly sampling the same population, or as a consequence of stratification due to the existence of different groups or conditions. In this setting, broadly five scenarios can be distinguished which are shown in the panels of Fig. 1. Fig. 1A represents the scenario where there is no variation across samples, with all samples in perfect equilibrium and having the same allele frequency. Fig. 1B shows three samples in perfect equilibrium but with different allele frequencies. In this situation, the variation in the genotype frequencies can be ascribed to variation in allele frequencies. Fig. 1C shows three samples with a different degree of deviation from equilibrium, but identical allele frequencies. In this scenario, the variation in the genotype frequencies is due to variation in the degree of disequilibrium and in ratio of homozygotes. Fig. 1D shows genotypic compositions that vary in allele frequency and in degree of disequilibrium, but that have a constant ratio of homozygotes. Finally, Fig. 1E shows samples that vary in allele frequency, in ratio of homozygotes and in degree of disequilibrium. Genetic polymorphisms may be regarded as doubly compositional because the division of the allele counts by their total establishes a second composition, the allelic composition. Some relationships between these two sets of compositions are established in this article (Section 2.3). In the remainder of this article, we provide some background on CoDA (Sections 2.1 and 2.2) and develop the logratio approach in the context of genetic equilibrium (Section 2.3). A multiallelic compositional measure for disequilibrium is presented, and its relationship with the classical inbreeding coefficient is shown. We explore the usefulness of the logratio approach for the analysis of polymorphisms using both simulations (Section 3) and practical genetic data analysis (Section 4) using biallelic data of the Glyoxalase locus (Ghosh, 1977; Olson, 1993) and multiallelic genotype data of STRs commonly used in forensics (Butler, 2006; Butler and Hill, 2012; Frontanilla et al., 2022).

## 2 Theory

This section gives a brief account of fundamental concepts in CoDA and of the exploration of compositional data by means of a logratio principal component analysis (LR-PCA), followed by the development of the logratio approach for the analysis of bi- and multiallelic polymorphisms.

### 2.1 Basic concepts of compositional data analysis

A composition of *D* parts is defined as a row vector of positive elements

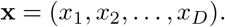

The sample space for a set of compositions is the simplex

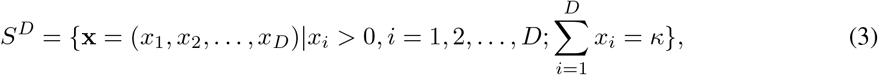

where *κ* > 0 is the closure constant. The closure operation *C* to the constant *κ* (usually *κ* = 1) is given by

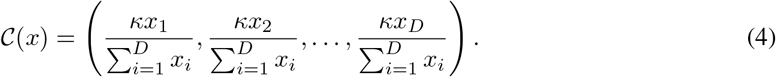

Notice that the formal definition of the simplex according to Eq. (3) as an *open simplex* (Aitchison, 1986) *excludes the zero outcome, whereas zero genotype counts are very common in genetics (see the Discussion section). For compositions, central tendency is measured by their geometric mean g*_*m*_(**x**), given by

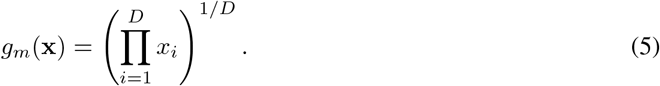

This corresponds to the ordinary mean in the logarithmic scale

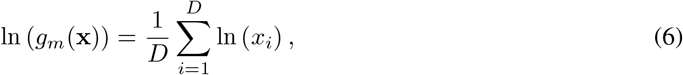

such that the geometric mean is easily obtained by exponentiation of the mean of the log-transformed data 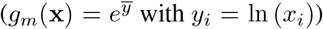 with *y*_*i*_ = ln (*x*_*i*_)). Dispersion in compositions is often expressed by the total compositional sample variance (Pawlowsky-Glahn et al., 2015), defined as

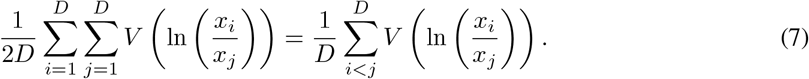

The mainstay of compositional data analysis is to free the compositional data from its unit-sum constraint by using logratio transformations. Currently, three different logratio transformations are in use. The additive logratio transformation (alr) concerns the logarithm of the ratios of two parts, where one part (not necessarily the last) is placed in the denominator and acts as a reference for all other parts

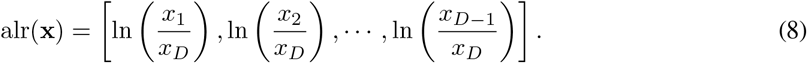

The centred logratio transformation (clr) concerns the ratios of one part against all parts in the form of their geometric mean

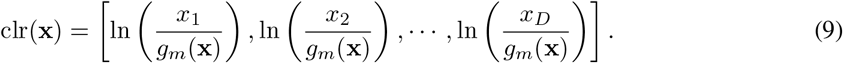

The isometric logratio transformation (ilr; Egozcue et al. (2003)) is given by

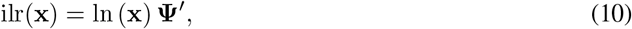

where **Ψ** is a *D −* 1 *× D* contrast matrix satisfying 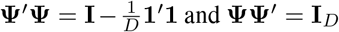 Isometric logratio coordinates encompass balances which are interpretable as ratios of geometric means of two subcompositions *i* and *j* composed of *r* and *s* parts respectively, and defined as

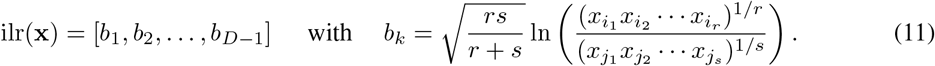

The contrast matrix **Ψ** corresponding to a balance can be constructed by a sequential binary partition (Pawlowsky-Glahn et al., 2015) of the parts of a composition; *r* and *s* are positive integers counting the number of positive (+1) and negative parts (*−*1) obtained in such a partition. The scalar 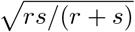 normalizes each row in the contrast matrix to unit sum-of-squares. We refer to Pawlowsky-Glahn et al. (2015) and Egozcue et al. (2003) for more details on constructing sequential binary partitions and contrast matrices for compositions. Examples of balances and their contrasts matrices in the current genetic context are found in Tables 1 and S1 below. Eq. (2) describes the HW law in logcontrast form; the general form of a logcontrast (Aitchison, 1986) is given by

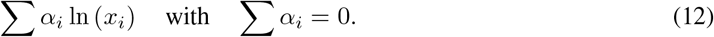

**Table 1:** Sequential binary partition and contrast matrix **Ψ** for a biallelic autosomal polymorphism.

| Seq. binary partition | | | | Contrast matrix $\Psi$ | | |
| --- | --- | --- | --- | --- | --- | --- |
| | $A_i A_i$ | $A_i A_j$ | $A_j A_j$ | | | |
| 1 | -1 | 1 | -1 | $-\frac{1}{2}\sqrt{\frac{2}{3}}$ | $\sqrt{\frac{2}{3}}$ | $-\frac{1}{2}\sqrt{\frac{2}{3}}$ |
| 2 | 1 | 0 | -1 | $\sqrt{\frac{1}{2}}$ | 0 | $-\sqrt{\frac{1}{2}}$ |

In this article, we will work with two genetic compositions, the allelic composition of a sample, and the genotypic composition of a sample. We define **p** to be the *K*-part allelic composition, and we will use **f** to denote the 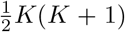-part genotypic composition, and use *p*_*i*_ and *f*_*i*_ to refer to the *i*th entries of these vectors respectively. Given that diploid genotypes consist of two alleles, the genotypic composition is also conveniently organised in the upper triangular *K × K* matrix **F**, and we use *f*_*ii*_ to refer to a diagonal (i.e., homozygote) element of **F** and *f*_*ij*_ to an above-diagonal (i.e., heterozygote) element. The order of the alleles in a heterozygote is considered irrelevant, such that the above-diagonal probabilities (with *i < j*) are 2*p*_*i*_*p*_*j*_ under HWE. We will also use **f** ^(*ii*)^ and **f** ^(*ij*)^ to subset the homozygotes and the heterozygotes of the full genotypic composition **f**.

### 2.2 Log-ratio principal component analysis

Aitchison (1983) developed LR-PCA for the exploration of compositional data, later further developed with compositional biplots (Aitchison and Greenacre, 2002). Here, we give a brief account of LR-PCA, where calculations are based on the singular value decomposition (SVD; (Eckart and Young, 1936)); this approach facilitates the construction of compositional biplots. Let **X**_*c*_ be the *n × D* compositional data matrix, and **X**_*l*_ = ln (**X**_*c*_) its elementwise logarithmic transformation. The column-centred matrix **X**_*c*clr_ of the clr transformed compositions is obtained by the double-centring operation **X**_*c*clr_ = **H**_*c*_**X**_*l*_**H**_*r*_, where **H**_*c*_ = **I** *−* (1*/n*)**11**^*′*^ and 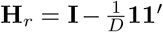 are the idempotent centring matrices for the columns and the rows of **X**_*l*_ respectively, after which the SVD follows:

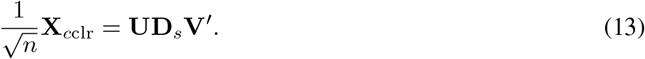

Biplot coordinates for the rows (**F**) and the columns (**G**) are calculated as

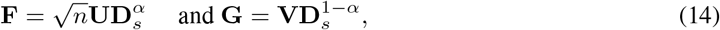

such that the clr transformed compositions are factored as **X**_*c*clr_ = **FG**^*′*^, and where *α* (0 ≤ *α* ≤ 1) is a conveniently chosen scalar. In practice, *α* = 1 and *α* = 0 are most often used, and the related coordinates are called standard (e.g., **F**_*s*_) an principal (e.g., **F**_*p*_) coordinates respectively, consequently given by

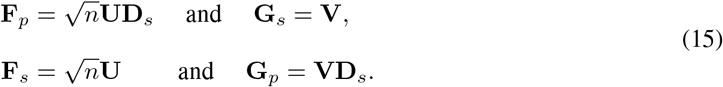

The factorisation **X**_*c*clr_ = **F**_*p*_**G**_*s*_^*′*^ is known as the *form* biplot, and **F**_*p*_ is the matrix with the principal components in its columns. Alternatively, a biplot based on the standardised principal components (**F**_*s*_) can be obtained as **X**_*c*clr_ = **F**_*s*_**G**_*p*_^*′*^, which is known as the *covariance* biplot. LR-PCA can also be carried out using the alr or ilr transformed compositions. The latter two types of analysis can be performed by applying the transformation, followed by column-centring and the SVD. Briefly, the corresponding matrices to be decomposed by the SVD are

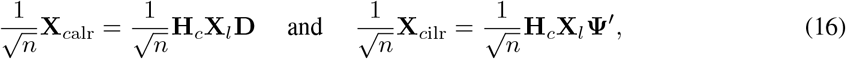

where **D** is a *D × D*(*D −* 1) difference matrix creating all pairwise differences of the columns of **X**_*l*_, and **Ψ** the aforementioned contrast matrix. In summary, there are three variants of LR-PCA depending on the logratio transformation that has been chosen. The three types of analysis are largely invariant, yielding eigenvalues and principal components that are identical up to a scalar. The main difference between the three types of analysis resides in the biplot vectors, which correspondingly either represent parts (clr), pairwise logratios (alr), or isometric logratios (ilr). The representation of supplementary information in biplots, either observations or variables, in biplots is possible by regression of conveniently transformed supplementary information onto the biplot axes (Dargie, 1984; Gabriel, 1995; Graffelman and Aluja-Banet, 2003). Doing this with compositional data has additional intricacies (Daunis-i Estadella et al., 2011).

### 2.3 The logratio approach to Hardy-Weinberg equilibrium

For a system with *K* alleles, *A*_1_, *A*_2_, … *A*_*K*_, there are 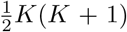 possible genotypes whose relative frequencies sum one. Let *f*_*ij*_ represent the genotype frequency of genotype *A*_*i*_*A*_*j*_ and let *p*_*i*_ represent the frequency of allele *A*_*i*_, such that 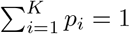. The transitive nature of the Hardy-Weinberg law (Graffelman and Weir, 2022) implies that for a particular population in HWE, the omission of all carriers of a certain allele from the population leads to a reduced genetic polymorphism that remains in HWE. Ultimately, if all alleles except *A*_*i*_ and *A*_*j*_ are eliminated, this implies Eq. (1) holds for each pair (*i, j*) of alleles such that

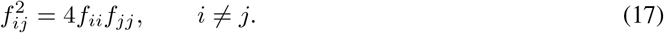

Consequently, beyond the unit-sum constraint ∑_*i*≤*j*_ *f*_*ij*_ = 1, there are 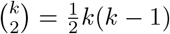 additional restric-tions on the *genotype space*, such that under equilibrium its dimensionality is given by

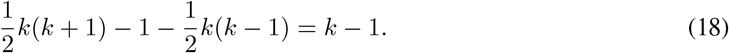

The number of alleles determines the dimensionality of the genotype space. The immediate consequence is that a biallelic genetic marker in equilibrium can be represented by an equilibrium line; the six genotype frequencies of triallelic marker can be perfectly represented in an equilibrium plane; and genotype compositions of multiallelic markers fall on an equilibrium hyperplane. Eq. (17) can be written in a ratio-based form, and by taking the square root we obtain

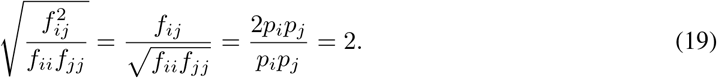

We note that Eq. (19) expresses that under equilibrium the ratio of the geometric mean of the heterozygote frequency and the geometric mean of the homozygote frequencies is the constant 2. When compared with Eq. (11) it appears directly proportional to the balance of heterozygotes and homozygotes. When the latter sare opposed in a sequential binary partition as shown in Table 1, and *A*_*i*_ alleles against *A*_*j*_ alleles in the second step, we obtain the balances

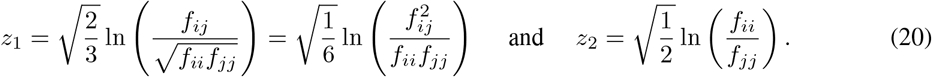

These balances have been previously studied in the biallelic setting (Graffelman and Egozcue, 2011). The following theorem generalises Eq. (19) to multiple alleles.

#### Theorem 1.

*Under HWE the geometric mean of the heterozygote frequencies of a polymorphism with K alleles is twice the geometric mean of the homozygote frequencies*.

*Proof*.

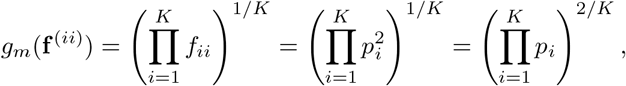

and

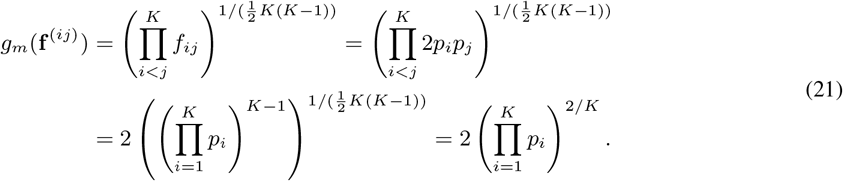

Upon taking ratios all allele frequencies cancel out such that

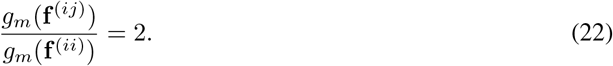

The geometric mean of the genotypic composition relates to the geometric mean of the allelic composition as shown next.

#### Theorem 2.

*Under HWE the geometric mean of a genotypic composition, g*_*m*_(**f**), *is the square of the geometric mean of its allelic composition, g*_*m*_ (**p**), *scaled by*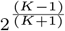.

*Proof*.

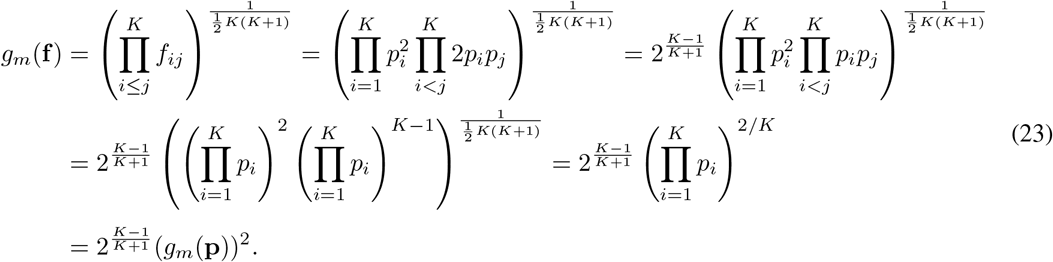

#### Theorem 3.

*Under HWE, the* clr *transformed parts of the allelic composition can be obtained by linearly combining the* clr *transformed parts of all corresponding carriers of the genotypic composition such that* 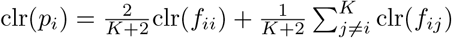

*Proof*. We have

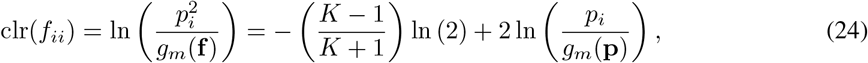

and

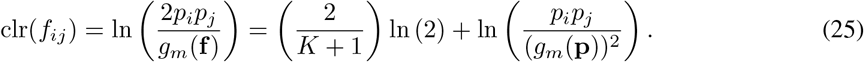

Now

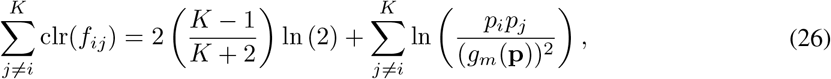

and

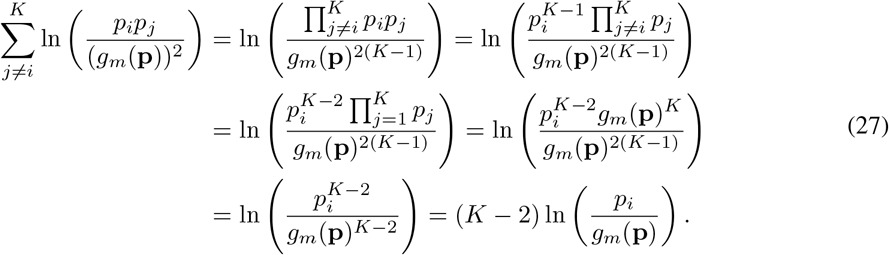

All terms involving ln (2) cancel out since

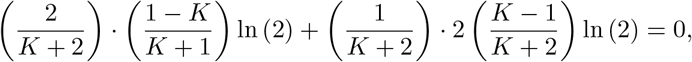

such that

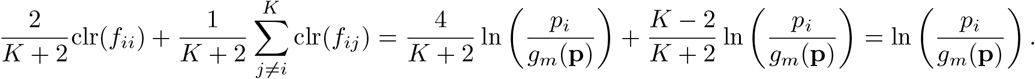

This result is useful for it allows one to represent the effects of the alleles in biplots of genotypic compositions under the HWE assumption (see Sections 3 and 4). Conversely, under HWE, the clr transformed parts of the genotypic composition are obtained by scalar multiplication or sums of clr transformed parts; this is immediately evident from (24) and (25) which can be written as

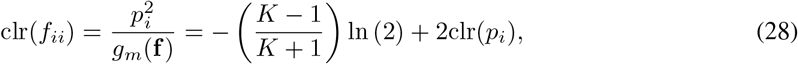

and

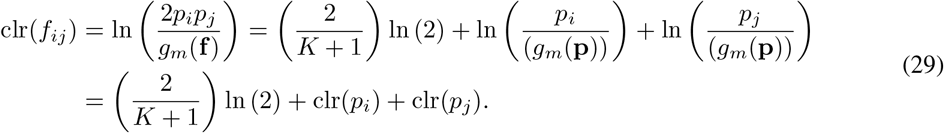

In PCA, the variables are column-centred, and consequently the effect of the leading scalars on the RHS of the latter two equations is removed. As a result, under exact HWE, a clr biplot vector of a homozygote is just twice the clr vector of its corresponding allele, and any heterozygote clr biplot vector is obtained by summing the clr vectors of its two alleles (see section 3.2). Similar to Theorem 2, the total compositional variance of the genotypic composition also has an exact relationship with the compositional variance of the allelic composition as shown below.

#### Theorem 4.

*Under HWE the total compositional variance of a genotypic composition, T*_*g*_, *of a polymorphism with K alleles is* (*K* + 2) *times the total compositional variance of its allelic composition, T*_*a*_.

*Proof*. The total compositional allelic variance is

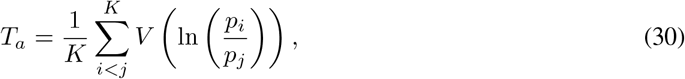

and the total compositional genotypic variance is

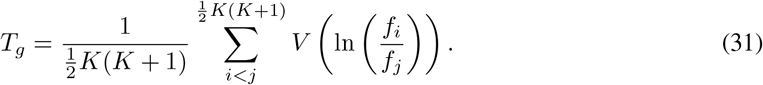

Both variances can be expressed as the sum of the variances of the clr transformed parts (Pawlowsky-Glahn et al., 2015), such that

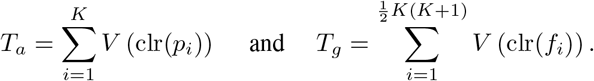

We split the latter sum into contributions due to the homozygotes 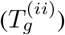 and the heterozygotes 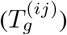, such that, under HWE, we have

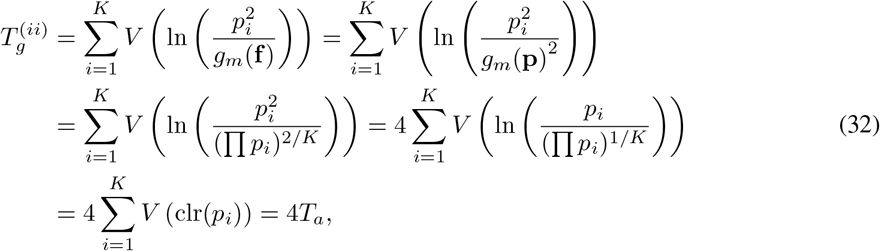

and

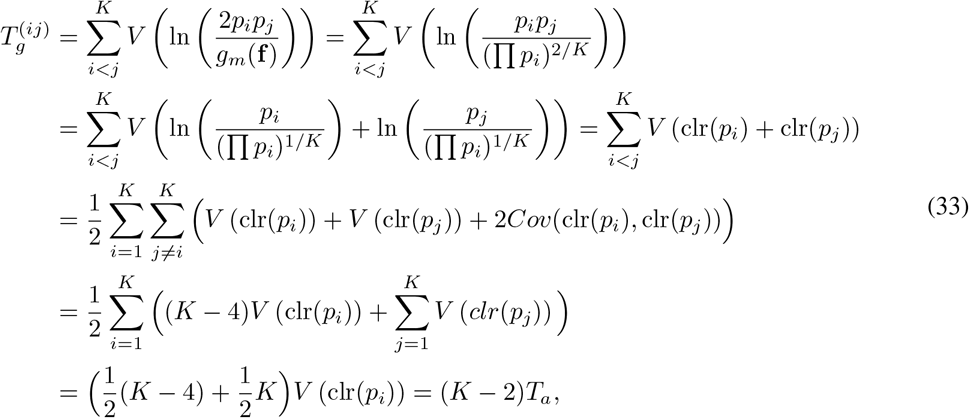

where we used Eq. (23) and the zero-sum constraint on the rows (and columns) of the covariance matrix of clr transformed compositions (Aitchison, 1986) given by

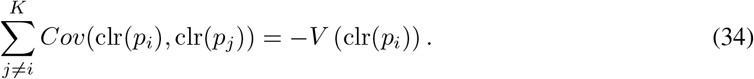

It thus follows that

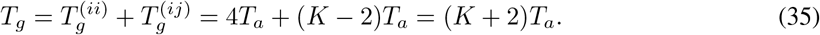

The latter equation holds true under exact HWE. Empirical data is subject to sample fluctuations which provoke the equation does not hold exactly for sample data, but is expected to hold approximately. An immediate consequence is that for a biallelic polymorphism under HWE, the total compositional variance of the genotypic composition is four times the variance of its allelic composition, or equivalently, twice the variance of the logit of its allele frequency.

The above theorems indicate that under HWE, the LR-PCA’s of the genotypic composition and the allelic composition are closely related. We use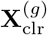 and 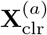 to refer to the clr transformed data matrix of genotypic and allelic compositions respectively, and use the same superscripts to refer to the results obtained from both types of analysis. From Eqs. (28) and (29) it is clear that all columns of 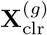 are linear combinations of the columns of 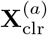, making that these matrices have identical rank. The relationship can be concisely expressed as

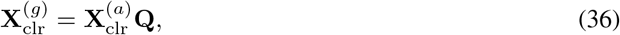

where **Q** is a 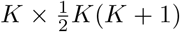 summing matrix that generates all pairwise sums of the columns of 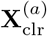, including sums of columns with themselves. The structure of **Q** is shown in Table 5 in the Appendix. Under exact HWE, the two LR-PCA’s are related in the following way:

- Both analyses have the same number of *K −* 1 non-zero eigenvalues. The analysis of the genotypic compositions has 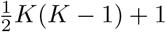 trailing zero eigenvalues and these dimensions can be ignored.
- The eigenvalues obtained in the two analyses satisfy *λ*^(*g*)^ = (*K* + 2)*λ*^(*a*)^. This result reaffirms Theorem 4. Consequently, the goodness-of-fit of a *k*-dimensional biplot of the genotypic and allelic compositions is the same, and the principal components obtained in both analyses are directly proportional, with coefficient of proportionality 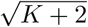.
- The eigenvectors, and consequently the biplot vectors obtained for the parts in both types of analysis, are related by a linear transformation given by 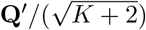. This result reaffirms Theorem 3.

These relationships are demonstrated in the Appendix.

Eq. (19) and its generalisation (22) naturally suggest that the ratio of the geometric means of heterozygotes over homozygotes can be used as a disequilibrium measure. For the biallelic case, the square of this ratio has been used in power calculations of tests for HWE (Rohlfs and Weir, 2008; Graffelman and Moreno, 2013). In epidemiology, half this ratio is known as *relative excess heterozygosity* (Ziegler et al., 2011). For a *K*-allelic polymorphism, the disequilibrium balance is given by

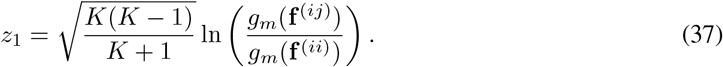

The dependence of this ratio on the scalar 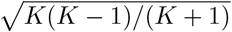 may, dependent on the context, be ignored, in which case the equilibrium reference value is ln (2), independently of the number of alleles. However, if a decomposition of the total genotypic variance is required, the scalar should be taken into account. The population inbreeding coefficient (*ρ*) is a commonly used measure for deviation from Hardy-Weinberg equilibrium in population genetics (Crow and Kimura, 1970; Hartl, 1980). In the biallelic scase, the genotype probabilities are modelled as 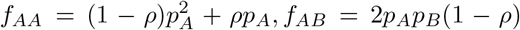 and 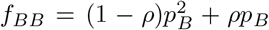 where *ρ* = 0 represents the equilibrium situation, *ρ <* 0 excess of heterozygotes and *ρ* > 0 a lack of heterozygotes. Parameter *ρ* can be estimated by maximum likelihood (Weir, 1996) as 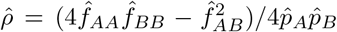. The term inbreeding coefficient may be considered inappropriate, for disequilibrium may derive from causes other than inbreeding. The mathematical relationship between the ratio-based measure of Eq. (17) and the population inbreeding coefficient has been described by Rohlfs and Weir (2008) for the biallelic case. For multiple alleles the genotype probabilities generalise as 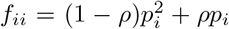 sand *f*_*ij*_ = 2*p*_*i*_ *p*_*j*_ (1 *− ρ*), and several estimators for *ρ* have been derived (Li and Horvitz, 1953). Figure 2 shows the relationship between *ρ* and the ratio of the two geometric means (see Eq. (22)) for different minor allele frequencies for the bi- and triallelic case; the compositional measure monotonously decreases in a non-linear fashion with *ρ*.

**Figure 2:**
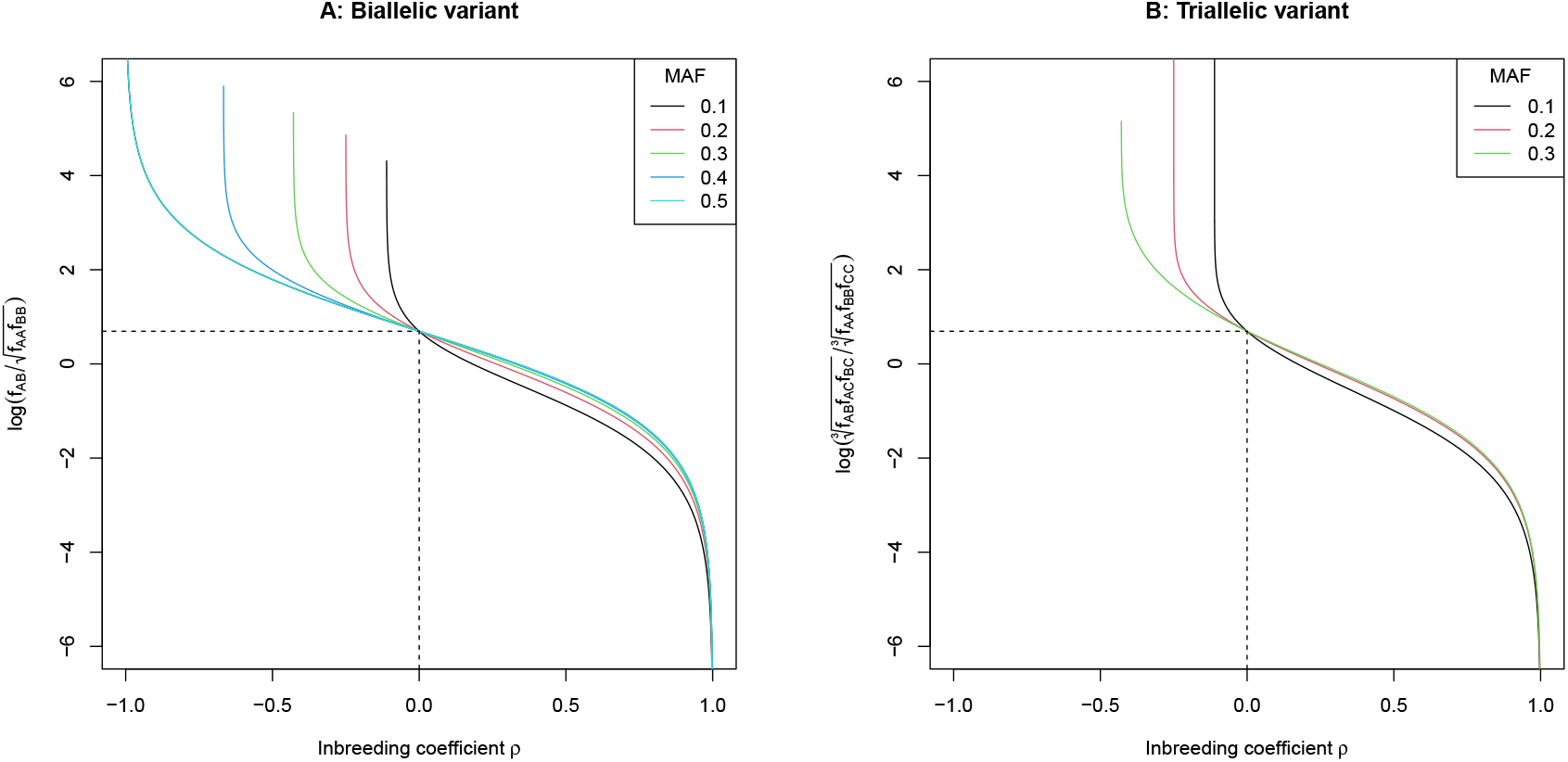
Relationship between the logratio of the geometric means of homozygotes and heterozygotes versus the inbreeding coefficient for different values of the minor allele frequency (MAF). A: biallelic polymorphism. B: triallelic polymorphism. The dashed line indicates the equilibrium values.

## 3 Simulations

In this section we apply LR-PCA to data simulated under HWE. Allele frequencies of *K*-allelic polymorphisms are drawn from a flat Dirichlet distribution with equal concentration parameters *Dir*(***α*** = **1**_*K*_). Genotype counts are obtained by taking samples of size 1,000 from a multinomial distribution with a parameter vector that equals the theoretical equilibrium genotype frequencies,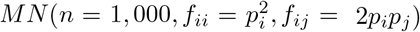. We intentionally use large sample sizes to reduce the amount of zero genotype counts and to illustrate the structures generated by HWE more clearly (see Discussion).

### 3.1 Biallelic polymorphisms

Figure 3A represents 1,000 samples of a biallelic polymorphism in a ternary diagram, and Figure 3B shows a biplot of the genotype compositions obtained by a clr covariance-based LR-PCA. The ternary diagram shows strong clustering of the samples around the Hardy-Weinberg parabola. In the LR-PCA biplot, the HW curve is represented by a horizontal line that coincides with the first principal component (i.e., *PC*_2_ = 0) which accounts for almost all (99.7%) of the compositional variance.

**Figure 3:**
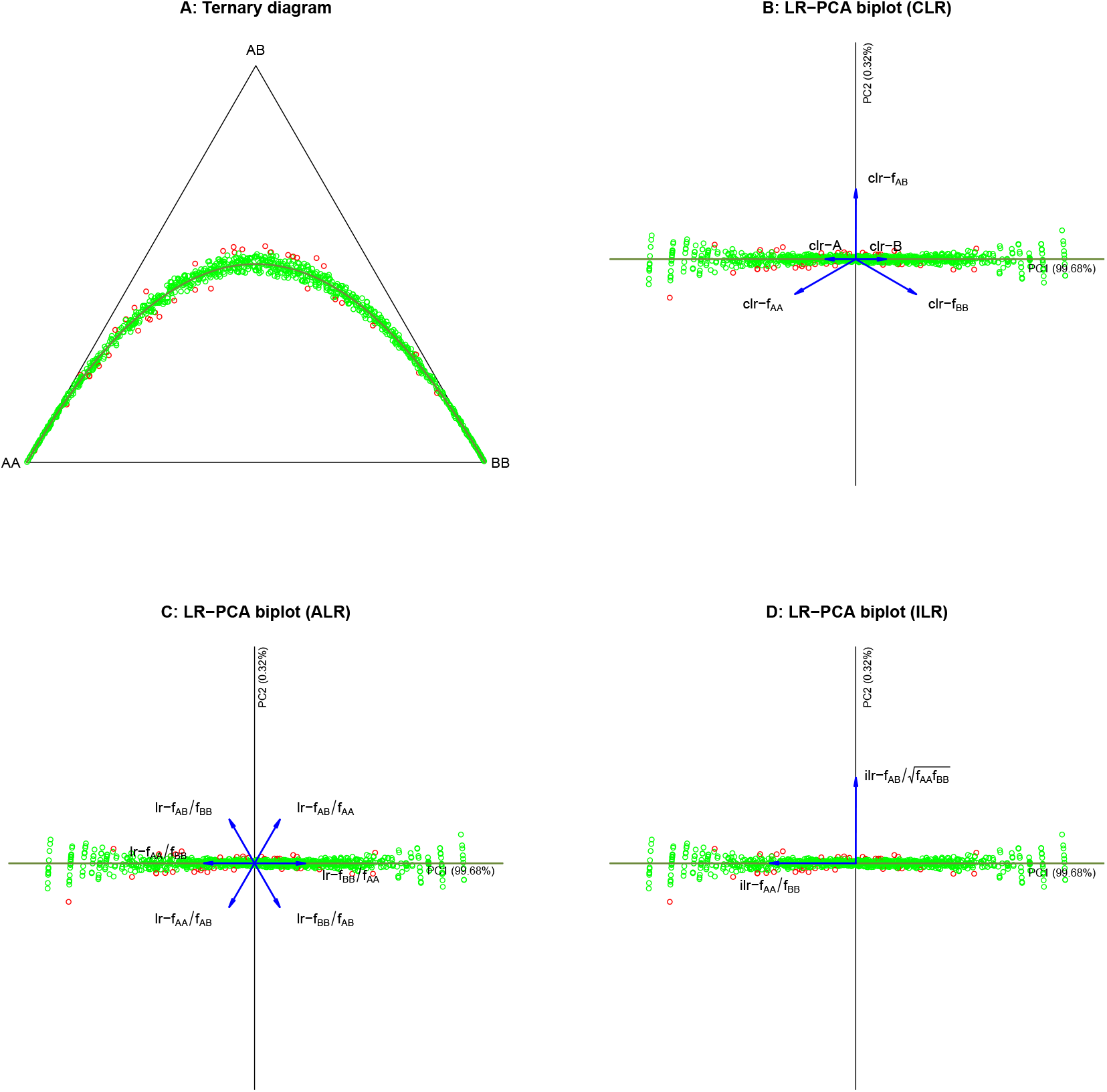
Ternary diagram and LR-PCA biplots of samples from a biallelic polymorphism. Biplot vectors are conveniently stretched by a constant to enhance the visualisation. A: Ternary diagram of 1,000 simulated genotype compositions with HW curve. B: The clr biplot of the genotypic compositions with equilibrium line. C: The alr biplot with all six pairwise logratios. D: The ilr biplot of the genotype compositions using the ilr transformation. Samples that are significant in an exact test for HWE are marked in red.

The variance decomposition and eigenvectors of the LR-PCA are given in Table 2. The first principal component is clearly interpretable as the log odds of the allele frequency, with eigenvectors coefficients that are close to 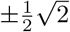, what, under HWE, leads to

**Table 2:** Variance decomposition and eigenvectors obtained in the LR-PCA of biallelic marker data.

| Variance decomposition |  |  |  | Eigenvectors |  |  |
| --- | --- | --- | --- | --- | --- | --- |
| | $PC_1$ | $PC_2$ | $PC_3$ | | $PC_1$ | $PC_2$ |
| $\lambda_i$ | 4.3409 | 0.0141 | 0.0000 | clr-AA | -0.7078 | 0.4071 |
| Fraction | 0.9968 | 0.0032 | 0.0000 | clr-AB | 0.0013 | -0.8165 |
| Cumulative | 0.9968 | 1.0000 | 1.0000 | clr-BB | 0.7064 | 0.4094 |

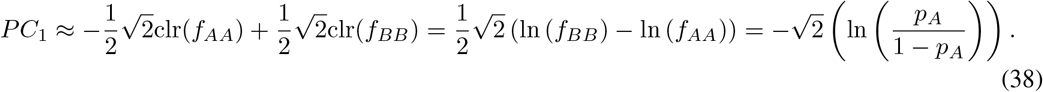

The eigenvalue of the last and third principal component is structurally zero due to the singularity of the clr covariance matrix; the second principal component is low-variance, and identifies a close-to-constant log-contrast:

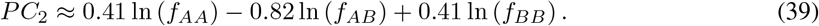

This logcontrast has a sample average of -0.56. It follows that

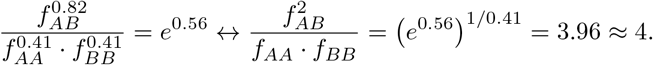

Clearly, the second principal component takes the logcontrast form of Eq. (2) and is a measure of Hardy-Weinberg disequilibrium. These interpretations of the components are easily confirmed by plotting the first and second component against the log odds of the sample allele frequency and the suggested disequilibrium measure respectively (see Figure S1). Applying Theorem 3, taking the linear combination of clr biplot vectors that carry the same allele, we obtain a biplot vector for the logit of the allele frequencies

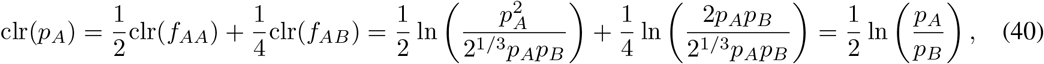

which is proportional to the first principal component; this way the clr and clr B vectors are represented in Fig 3B. Fig 3C shows the biplot of the alr based LR-PCA which directly shows all pairwise logratios; logratio *f*_*AA*_*/f*_*BB*_ directly identifies *PC*_1_ as the log odds of allele A, assuming HWE. The logratios *f*_*AB*_*/f*_*AA*_ and *f*_*AB*_*/f*_*BB*_ sum to the second principal component, which is thus also identified as the disequilibrium measure 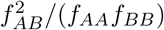. Figure 3D shows a LR-PCA biplot obtained by using the ilr transformation. The PCs clearly coincide with the isometric logratios *z*_2_ and *z*_1_ respectively. In summary, under HWE, the LR-PCA of samples of an autosomal biallelic genotype composition can uncover two interpretable, orthogonal dimensions, one driven by the variation in allele frequency as expressed by its logit, and one driven by a disequilibrium measure, the balance between heterozygotes and homozygotes. The significant samples are, for the given value of the log odds of the allele frequency, seen to be most outlying with respect to the equilibrium line.

### 3.2 Multiallelic polymorphisms

1,000 triallelic markers were simulated using multinomial samples of size *n* = 1, 000 with Hardy-Weinberg probabilities and allele frequencies drawn from a *Dir*(1, 1, 1) distribution. Figure 4A shows the two-dimensional LR-PCA biplot of the genotypic compositions, where samples with a significant exact test for HWE (*α* = 0.05) are coloured in red. Figure 4A reveals a profound structure in the data. First of all, the biplot is an almost perfect fit of the genotypic compositions in two dimensions, as all eigenvalues beyond the second are almost zero (see the sharp drop in the screeplot in Fig. S2), as is also predicted by Eq. (18). Secondly, three triples of collinear biplot vectors appear, as marked by the dashed lines in Figure 4A. Each set of collinear vectors involves only two alleles. These collinear vectors are indicative for a one-dimensional relationship for the corresponding biallelic subcompositions. When these subcompositions are actually calculated and analysed, the aforementioned structure for a biallelic polymorphism as exposed in the previous subsection appears (see Figure S3). Figure 4A represents the *equilibrium plane* and shows the *nested* nature of HWE, as the equilibrium plane contains three equilibrium lines, one for each biallelified polymorphism. Finally, Figure 4 shows 12 orthogonal logratios, where the bisector of each homozygote vertex (e.g., CC-AB) is orthogonal to the logratio of the heterozygotes both carrying the homozygote’s allele (e.g., AC and BC), and orthogonal to the logratio of the other two homozygotes (e.g. AA-BB). The aforementioned collinearity triples implies the bisector is also orthogonal to the heterozygote-homozygote logratios (e.g. AB-AA and AB-BB). The three alleles of the clr transformed allelic compositions are represented in Figure 4A by using Theorem 3. In this case, the vectors representing the alleles are, as expected, proportional to their corresponding homozygote vectors. The allele vectors clearly show the first principal component can be interpreted as the log odds of the B and C allele frequencies, whereas the second principal component is the balance of the geometric mean of the A allele over the geometric mean of the B and C alleles. These interpretations are verified when the principal components are plotted against the aforementioned quantities (see Fig. S4). Figure 4B shows the plane of the first two principal components, and the third principal component as the orthogonal vertical dimension. The eigenvectors of the analysis in Table 3 show that the third component clearly opposes homozygotes to heterozygotes and comes close to being the ratio of the geometric means of homozygotes and heterozygotes. Compositions above the plane (light blue) have an excess of homozygotes; compositions below the plane (dark blue) an excess of heterozygotes. For triallelic variants, an ilr based LR-PCA can be performed with the heterozygote-homozygote contrast as the natural first contrast, followed by pivot-like within-homozygote and within-heterozygote contrasts (see Table S1). Since the variance decomposition is the same for both types of analysis, it is not surprising that the third component of the latter analysis indeed identifies the heterozygote-homozygote balance (see the first line in Table S2).

**Table 3:**
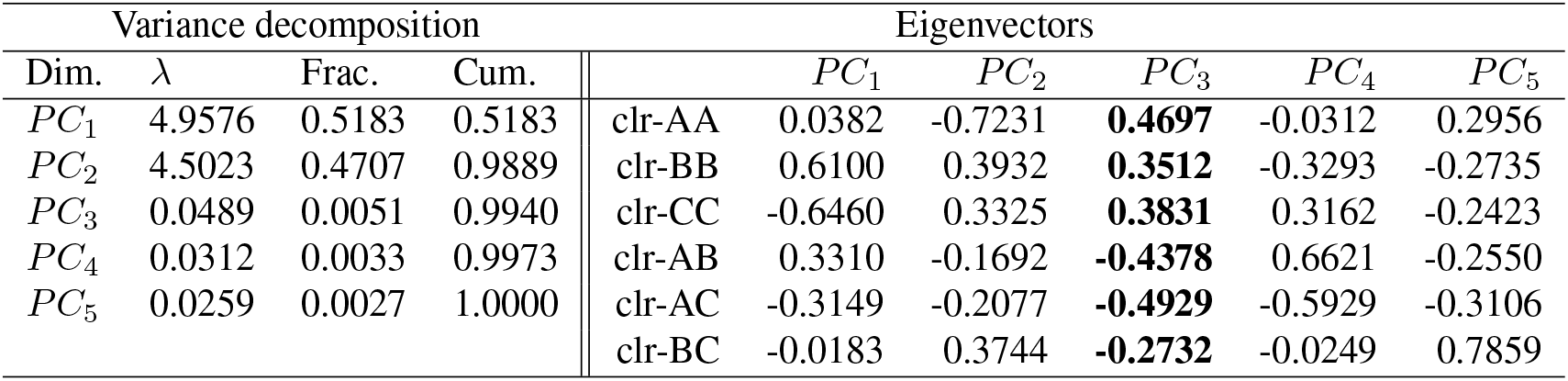
Variance decomposition and eigenvectors obtained in the clr based LR-PCA of multiple samples from a triallelic polymorphism under HWE. Coefficients of *PC*_3_ (in bold) show its interpretation as a homozygote-heterozygote balance.

| Variance decomposition |  |  |  | Eigenvectors |  |  |  |  |  |
| --- | --- | --- | --- | --- | --- | --- | --- | --- | --- |
| Dim. | $\lambda$ | Frac. | Cum. | | $PC_1$ | $PC_2$ | $PC_3$ | $PC_4$ | $PC_5$ |
| $PC_1$ | 4.9576 | 0.5183 | 0.5183 | clr-AA | 0.0382 | -0.7231 | **0.4697** | -0.0312 | 0.2956 |
| $PC_2$ | 4.5023 | 0.4707 | 0.9889 | clr-BB | 0.6100 | 0.3932 | **0.3512** | -0.3293 | -0.2735 |
| $PC_3$ | 0.0489 | 0.0051 | 0.9940 | clr-CC | -0.6460 | 0.3325 | **0.3831** | 0.3162 | -0.2423 |
| $PC_4$ | 0.0312 | 0.0033 | 0.9973 | clr-AB | 0.3310 | -0.1692 | **-0.4378** | 0.6621 | -0.2550 |
| $PC_5$ | 0.0259 | 0.0027 | 1.0000 | clr-AC | -0.3149 | -0.2077 | **-0.4929** | -0.5929 | -0.3106 |
|  |  |  |  | clr-BC | -0.0183 | 0.3744 | **-0.2732** | -0.0249 | 0.7859 |

**Figure 4:**
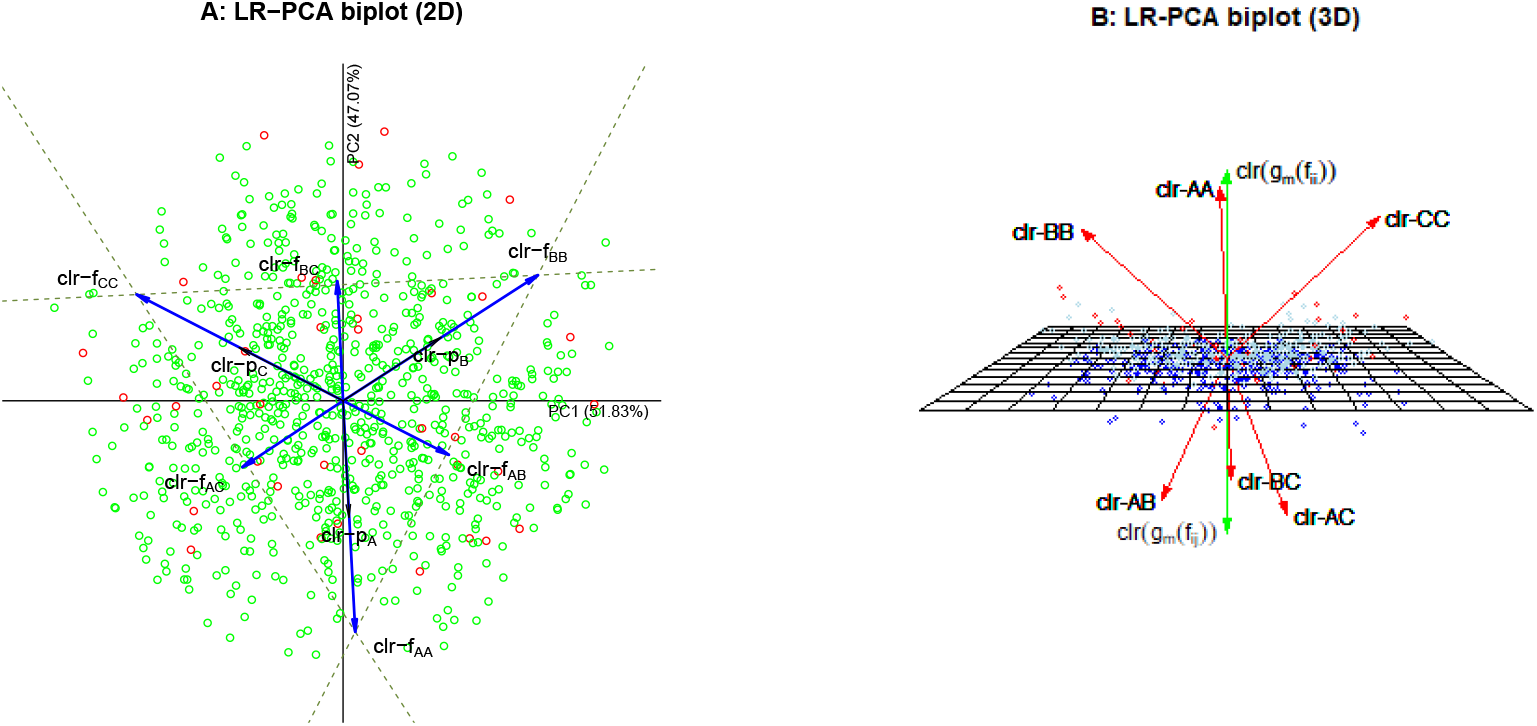
A: LR-PCA biplot of the first two principal components of 1,000 simulated samples from a triallelic genetic marker under Hardy-Weinberg equilibrium. B: Three dimensional plot of the first three principal components with projections onto the plane spanned by the first two components. The two green vectors represent the geometric averages of homozygotes and of the heterozygotes. Samples significant (*α* = 0.05) in an exact test for HWE are marked in red.

We increase the dimensionality and simulate another 1,000 samples, now of a five-allelic polymorphism, both under exact equilibrium (exact HWE probabilities) and statistical equilibrium (multinomial sampling with HWE probabilities). The LR-PCA biplots using the clr transformation are shown in Figure 5. Though the exact biplots (Figs. 5A, 5B) account for only 53.2% of the compositional variance, the pattern of biplot vectors is highly informative. All triples of biplot vectors involving a single pair of alleles appear exactly collinear. This is illustrated in Fig. 5A for the pairs (A,B), (A,C), (A,D) and (A,E) with dashed lines, but it holds exactly for all ten pairs. Fig. 5B shows the biplot of the LR-PCA of the allelic compositions, and shows how the homozygote vectors AA and BB and the heterozygote vector AB are obtained by doubling and summing the corresponding clr(*p*_*A*_) and clr(*p*_*B*_) vectors, according to Theorem 3. Doing so exactly reproduces the configuration of the genotypic vectors in Fig. 5A. Biplot vectors in Fig. 5B are scaled to match those of Fig. 5A, but the principal components are left unscaled. This clearly shows the increased variance of the components obtained in the LR-PCA of the genotypic compositions. Figs. 5C and 5D show the analysis of genotypic compositions, but for data generated from a multinomial distribution assuming known allele frequencies and equilibrium probabilities for two different sample sizes, 1,000 and 10,000 respectively. The first setting generated 5.5% of zero genotype counts, which were imputed using a multiplicative count zero replacement (Palarea-Albaladejo and Martín-Fernández, 2015). Though the overall percentage of zero counts is low, due to the large size of the compositions (15 genotypes), 45% of the observations (i.e., rows) in the data matrix has at least one zero. Due to the noise generated by multinomial sampling and zero imputation, most of the expected collinearity patterns appear broken in Fig. 5C, where the heterozygote vectors are not on the line segments joining their corresponding homozygotes. When allelic vectors are added, using the HWE assumption, clr vectors still appear proportional to their corresponding homozygote vectors. Samples with a larger number of zeros tend to be peripheral in the plot. If the counts of the simulated samples are tested for HWE (at *α* = 0.05), 5.2% is significant, close to what is expected by chance alone. If, after imputation of the zeros, the data is scaled back to count data, the rejection rate remains 5.2%. The screeplot of the eigenvalues (see Fig. S5) reveals a sharp drop from the fourth to the fifth eigenvalue, which is expected as the theoretical dimensionality under HWE is four (see Eq. 18). The eigenvectors (Table S3) shows the fifth eigenvector can be interpreted as a heterozygote-homozygote balance, having negative coefficients for all heterozygotes and positive coefficients for all homozygotes, with the two sets of coefficients having fairly similar within-set magnitudes. This interpretation of the fifth component is confirmed by plotting it against the heterozygote-homozygote balance (see Fig. S6). Fig. 5D shows the same analysis but for the larger samples size of *n* = 10, 000, which has 1.5% zero counts with 17% of the observations affected by the need for zero-imputation. At this larger sample size the expected collinearity patterns are close to being perfectly restored. The statistical testing of the original multinomial samples (i.e, with the zeros) and the zero-imputed samples is similarly calibrated (3.7% rejections in both cases). The LR-PCA’s of the allelic compositions corresponding to Figs. 5C and 5D (not shown) have a slighty different goodness-of-fit statistics in comparison with the genotypic LR-PCA, which is expected since there is no exact equilibirum such that Eq. (46) does not hold exactly.

**Figure 5:**
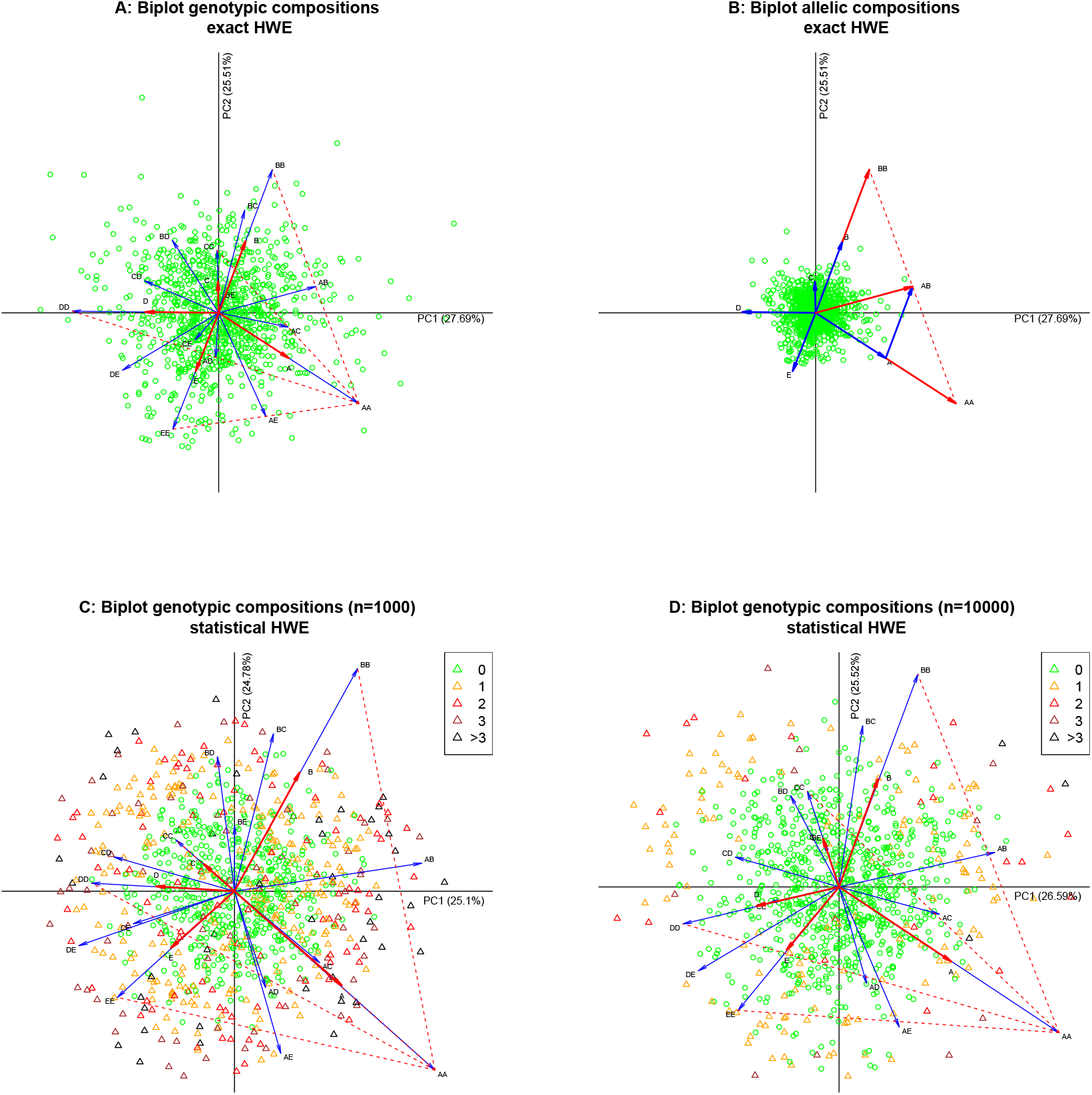
LR-PCA biplot of the first two principal components of 1.000 simulated samples from a five-allelic genetic marker under exact Hardy-Weinberg equilibrium (panels A and B) and statistical equilibrium (panels C and D). A: biplot of genotypic compositions with added allelic vectors in red. In this panel biplot vectors are scaled by a factor of 20 to enhance the visualisation. B: biplot of allelic compositions with added genotypic vectors (in red) for genotypes AA, BB and AB. In panel B, allelic biplot vectors are scaled by 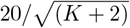 to match panel A. C: Biplot of genotypic compositions resulting from multinomial sampling (*n* = 1, 000) and zero imputation, with added allelic vectors in red. D: Biplot of allelic compositions resulting from multinomial sampling (*n* = 10, 000) and zero imputation. Some collinearities are shown by dashed lines connecting homozygotes. The prefix “clr” is omitted from the biplot vector labels to facilitate the visualisation. Samples with one or more zeros are marked with a triangle, and coloured according to the number of zeros.

## 4 Empirical genetic data analysis

We apply the methodology hitherto exposed to a biallelic data from the glyoxalase locus (Ghosh, 1977; Olson, 1993) and to multiallelic STR data derived from the 1,000 genomes project (The 1000 Genomes Project Consortium, 2015; Frontanilla et al., 2022).

### 4.1 Biallelic polymorphisms

We analyze glyoxalase genotype data for 17 populations from India (Ghosh, 1977; Olson, 1993; Olson and Foley, 1996). A ternary diagram of the samples and the LR-PCA biplot are shown in Figures 6A and 6B. For these populations, the B allele is the major allele. When each population is tested separately for HWE with an exact test, equilibrium is rejected for one population (Naicker). Figure 6B shows the biplot of a clr based LR-PCA. An equilibrium line has been added to the biplot, and is obtained by adding genotype compositions consisting of the expected genotype frequencies under HWE to the plot as supplementary points. Quite contrary to the results obtained by simulation (see Fig. 3), the equilibrium line does not coincide with the first principal component, and for this data, the second principal component accounts for 22.2% of the compositional variance, and is not low-variance. The eigenvectors obtained in this analysis are given in Table 4 and, unlike the vectors obtained in the simulations (see Table 2), do not evidentiate the HW law. When LR-PCA is applied using the ilr transformation, the compositional biplot in Figure 3C is obtained. All the different versions of LR-PCA yield, as expected, exactly the same variance decomposition. The ilr biplot shows the same configuration of the populations, with two orthogonal biplot vectors representing the isometric logratios *z*_2_ = ln (*f*_*AA*_*/f*_*BB*_) and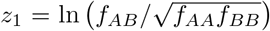. Panels 6B and 6C appear rotated with respect to the structures observed in the simulations (see Fig. 3). Biplot vector ln (*f*_*AA*_*/f*_*BB*_) coincides with the equilibrium line; biplot vector ln 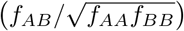is orthogonal to the equilibrium line and represents deviation from equilibrium. The principal components in biplots in panels 6B and 6C are hard to interpret, they have no obvious genetic meaning. The directions provided by the ratios ln 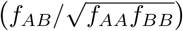and ln (*f*_*AA*_*/f*_*BB*_) are genetically the most interpretable. Consequently, the decomposition of genotypic compositional variance over the principal components is of less interest; a different decomposition of this variance over the two ilr’s is of more interest and has a clear genetic interpretation. This decomposition is shown in the last two columns of Table 4, it can thus be inferred that 65% of the observed genotypic variance can be ascribed to variation in the homozygote ratio across the samples, and 35% is due to variation in the disequilibrium measure. The clr and ilr biplots in panels 6B and 6C may be rotated in order to render the components more interpretable such that they coincide, as previously observed in the simulations, with the homozygote ratio and the disequilibrium measure. However, such rotation is fact not needed for one can just calculate the ilr coordinates directly from the genotype frequencies and plot them on the orthogonal axes of an ordinary scatterplot as shown in Figure 6D to obtain the desired simplification.

**Table 4:** Decomposition of genotypic compositional variance for the Glyoxalase data. The first six columns give the variance decomposition and eigenvectors obtained in a LR-PCA using the clr transformation. Columns seven through nine give the eigenvectors obtained in LR-PCA with the ilr transformation. The last two columns given the decomposition of the compositional variance using plain ilr coordinates without performing LR-PCA.

| LR-PCA |  |  |  |  |  |  |  |  | Scatterplot |  |
| --- | --- | --- | --- | --- | --- | --- | --- | --- | --- | --- |
| Var. decomposition |  |  | Eigenvectors |  |  |  |  |  | Var. decomposition |  |
|  |  |  | clr based |  |  | ilr based |  |  |  |  |
| | $PC_1$ | $PC_2$ | | $PC_1$ | $PC_2$ | | $PC_1$ | $PC_2$ | $z_1$ | $z_2$ |
| $\lambda$ | 0.2340 | 0.0666 | clr-AA | -0.8164 | -0.0107 | $z_1$ | 0.4886 | 0.8725 | 0.1133 | 0.2062 |
| Fraction | 0.7784 | 0.2216 | clr-AB | 0.3989 | 0.7124 | $z_2$ | -0.8725 | 0.4886 | 0.3545 | 0.6455 |
| Cumulative | 0.7784 | 1.0000 | clr-BB | 0.4175 | -0.7017 |  |  |  | 0.3545 | 1.0000 |

**Figure 6:**
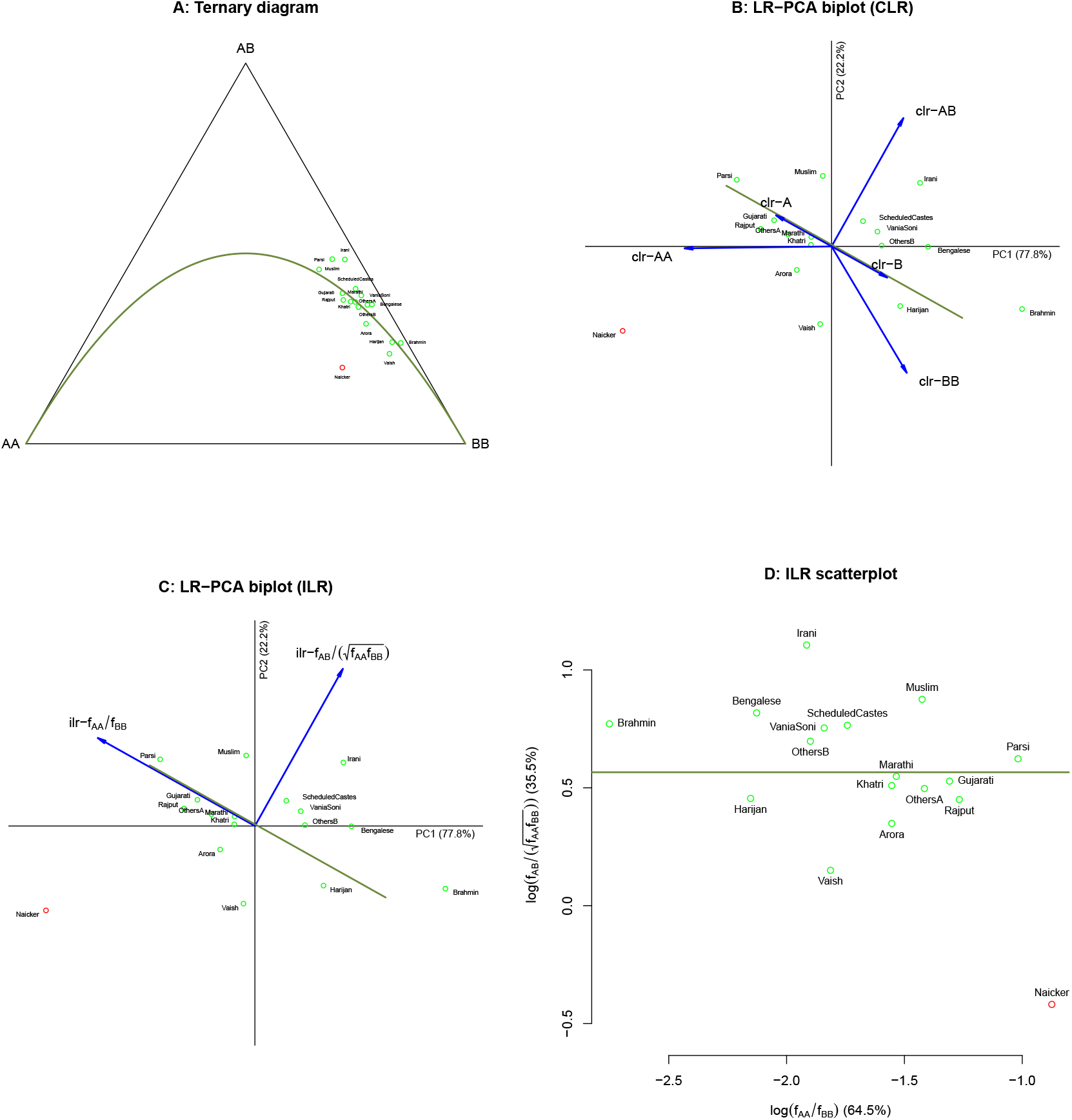
Glyoxalase genotype data for 17 populations from India. Populations for which HWE is rejected are marked in red. The HWE equilibrium condition is indicated with an olive-green curve or line. A: Ternary diagram of the genotype frequencies with HW parabola. B: LR-PCA biplot based on the clr transformation, with equilibrium line. C: LR-PCA biplot based on the ilr transformation. D: Scatterplot of the ilr transformed genotype frequencies. A reference equilibrium line is shown at 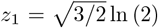 ln (2).

### 4.2 Multiallelic polymorphisms

Frontanilla et al. (2022) generated 21 STRs for the 2405 individuals of the 26 populations of Phase 3 of the 1,000 Genomes project (The 1000 Genomes Project Consortium, 2015); its data is available at www.internationalgenome.org. To analyse a particular *K*-allelic STR by LR-PCA, the 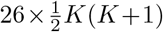 matrix of genotype counts was calculated. Because STRs have many alleles, many of them with low frequency, this matrix it typically highly sparse. Sparseness was reduced by merging alleles, which theoretically retains equilibrium, if present (Graffelman and Weir, 2022). Typically, the shortest and the longest alleles are less frequent, and STRs were reduced to three or four alleles by joining shorter alleles and longer alleles with the more common adjacent alleles. Fractional alleles were joined with their corresponding non-fractional allele. This procedure typically produces STRs that have no or just a few zeros. Any remaining zeros were imputed by using a multiplicative count zero replacement (Palarea-Albaladejo and Martín-Fernández, 2015). We describe the analysis of D2S441. This STR was reduced to triallelic by aggregating the tails of the allele frequency distribution, such that three alleles remain (≤ 10, 11, ≥ 12), reducing the number of zero genotype counts to 4.5%.

Figure 7A shows the LR-PCA biplot of D2S441. There is a clear clustering of populations according to their population grouping, with AFR populations on the left and SAS on the right of the plot. AMR populations which are known to be admixed, appear blurred. The African-American Southwest US population (ASW) and the African-Caribbean population (ACB) cluster are close to, as might be expected, the AFR populations. Biplot vectors for homozygote genotypes are joined by line segments, which show that biplot vectors pertaining to each of the pairs (10,11), (10,12) and (11,12) are approximately collinear, as would be expected under HWE. The total compositional genotypic variance of the reduced triallelic STR is 3.2535 of which only 2.4% can be attributed to disequilibrium measure *z*_1_. The analysis is repeated without the admixed AMR samples in Figures 7C and 7D, where the four population groups are almost perfectly separated. The LR-PCA’s in Fig. 7 have an expected dimensionality of two; the disequilibrium measure *z*_1_ is seen to correlate with the third principal component in both cases (see Fig. S7).

**Figure 7:**
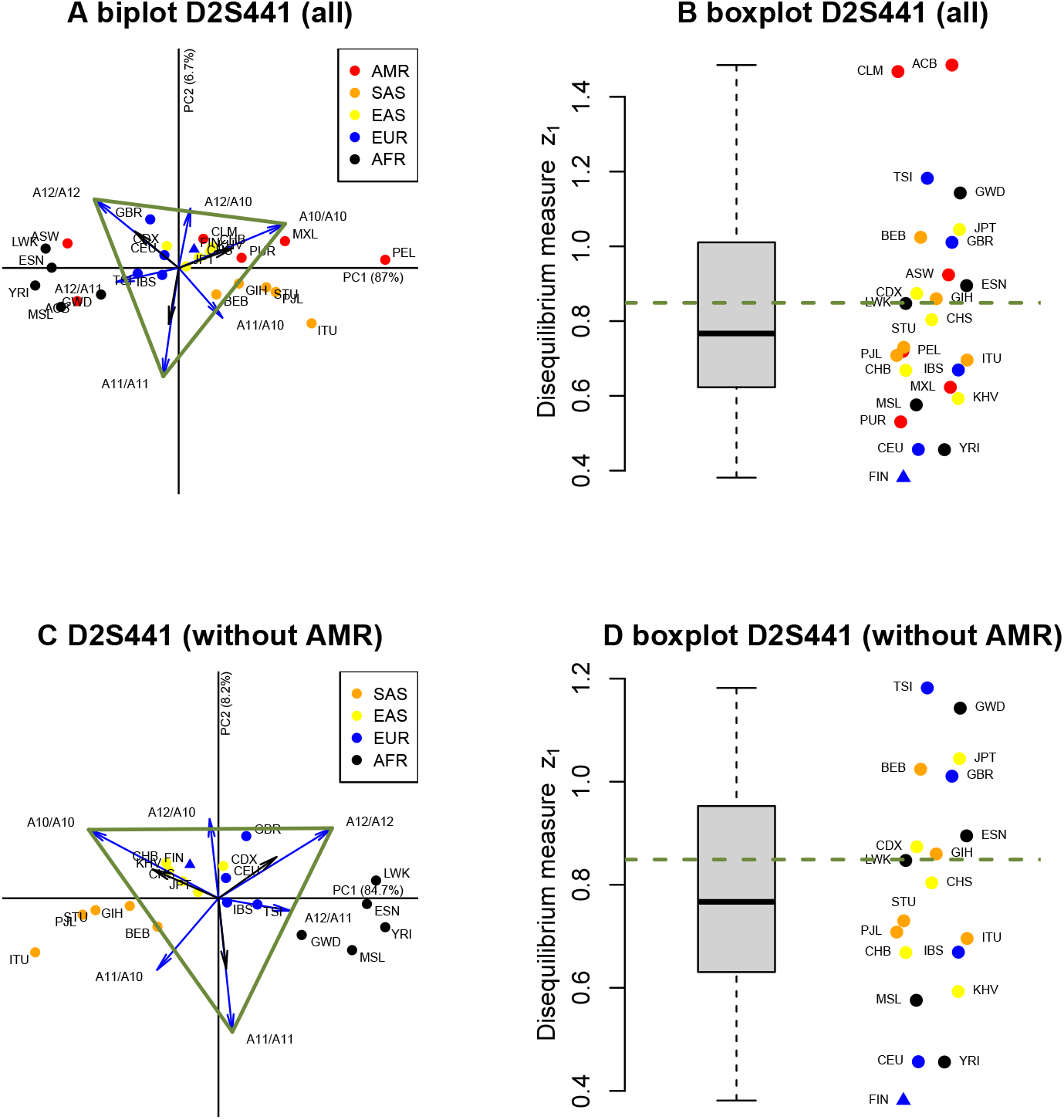
Biplots and disequilibrium boxplots of triallelified STR D2S441. Colouring of dots according to the population groups African (AFR), admixed American (AMR), East Asian (EAS), European (EUR) and South Asian (SAS). The three-letter population coding is according to the 1000G project. LR-PCA biplot vectors are stretched by a factor of three to facilitate the visualisation. Homozygotes are connected by line segments to show approximate collinearity for reduced biallelic polymorphisms. Samples significant in an sexact test for HWE (*α* = 0.05) are marked with a triangle. A: LR-PCA biplot. B: boxplot of disequilibrium measure *z*_1_. C: LR-PCA biplot of LR-PCA without AMR samples. D: boxplot of disequilibrium measure *z*_1_ without AMR samples. The equilibrium reference value of 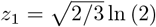, is indicated in the boxplots by a dashed horizontal line.

## 5 Discussion

In genetic data analysis, genotype counts are commonly modelled with a multinomial distribution whose probability vector is determined by the HW law. Zero genotype counts are very common, and are expected in particular for homozygotes of rare alleles. In compositional data analysis compositions are usually defined as consisting of positive elements only (see Eq. (3)). There is an extensive literature on dealing with zeros in compositions (Fry et al., 2000; Martin-Fernandez et al., 2003; Palarea-Albaladejo and Martín-Fernández, 2015). The fact that polymorphism data is typically fraught with zeros has hampered the application of the logratio approach in genetics. Nowadays, in genetic association studies very large sample sizes are increasingly being used, with sample sizes running up to half a million in the UK biobank project (Bycroft et al., 2018). Such large samples reduce the number of zeros and make the logratio approach more feasible. The merging of alleles can also greatly alleviate the zero problem as shown in the multi-allelic example, though it has several drawbacks. For STRs, disequilibrium is often due to rare homozygotes, which are highly unlikely under HWE (Graffelman and Weir, 2022). Merging the rare alleles with more common alleles converts these rare homozygotes into more common homozygotes, and thus brings about a loss in power to detect disequilibrium. Likewise, if there are groups in the data (i.e., population substructure or treatment groups) the fusion of alleles may obliterate group differences and diminish power to detect group effects.

The LR-PCA of genotype frequencies has shown to be informative, for it can simultaneously visualise ratios of genotype and allele frequencies and it is informative about HWE. However, even in the biallelic case inferring HWE is more complex than apparent in the work of Aitchison (1999). The analysis of the Glyoxalase data in Figure 7 shows the information regarding HWE is not necessarily a direction of minimum variance, and is therefore neither necessarily represented by the second principal component. Ultimately, for a biallelic variant, it is shown that LR-PCA can be circumvented, and that all variation in the genotypic compositions can be efficiently presented in single scatterplot of two logratios, which allows for a genetically more meaningful breakdown of the total genotypic variance. LR-PCA is nevertheless recommended, notably so for multiallelic variants, because it allows for the construction of compositional biplots. The compositional biplots enables one to spot equilibrium collinearity patterns which are hard to infer by merely looking at the coefficients of the eigenvectors (see Figures 4, 5, 7). The compositional biplot may suggest certain biallelifications that are likely to satisfy the equilibrium assumption, even when equilibrium is not met overall. Moreover, these biplots are very useful for detecting outliers and clusters of samples. The LR-PCA of the genotype frequencies has, under HWE, a theoretical dimensionality *K −* 1, and therefore a drop in the eigenvalues towards zero at *K* is expected (see Figs. S2, S5), though not always observed in practice. Simulations and practical data analysis show the heterozygote-homozygote balance *z*_1_ (Eq. (37)) tends to appear in LR-PCA. However, and importantly, LR-PCA at best approximates this measure. The separate, direct calculation of this balance from the genotype frequencies and its inspection (see Figs. 7B and 7D) is therefore recommended.

The theorems derived in Section 2.3 are largely confirmed by the matrix equations in the Appendix. The theorems as such do not consider the low-dimensional approximation of logratio transformed allelic and genotypic compositions. The results in the Appendix show the relations between allelic and genotypic logratio transformed parts do equally apply for low-rank approximations in a low-dimensional subspace. This is most clearly illustrated in Figures 5A and 5B which are far from perfect approximations to the higher-dimensional compositions, but where all rules for obtaining allelic vectors from genotypic vectors or the reverse, validly apply. This ultimately means that the collinearity between the triple of biplot vectors of the homozygotes *A*_*i*_*A*_*i*_, *A*_*j*_*A*_*j*_ and their heterozygote *A*_*i*_*A*_*j*_ is, under exact HWE, in principle *always* expected to occur irrespective of the number of alleles of the polymorphism. This may be surprising at first sight. Whenever this collinearity is *not* observed, it is not explained by error in the low dimensional approximation to the high-dimensional allelic and genotypic composition, but it is due to some degree of deviation from Hardy-Weinberg equilibrium. This occurs in, for instance, Figures 3B, 6B and 7C. Importantly, perfect collinearity is expected under exact HWE; whenever missing data or zeros are imputed, this will likely generate some degree of disequilibrium, and consequently deviations from perfect collinearity.

## 6 Acknowledgements

This work was supported by the Spanish Ministry of Science and Innovation and the European Regional Development Fund under grant PID2021-125380OB-I00 (MCIN/AEI/FEDER).

The author reports there are no competing interests to declare.

## Appendix Relation between the LR-PCA’s of the allelic and the genotypic compositions

Table 5 gives matrix **Q** for a four-part composition.

Note the columns of **Q** all sum to two and its rows sum to *K* + 1. Inner products between the rows of **Q** equal 1. Consequently, we have

**Table 5:**
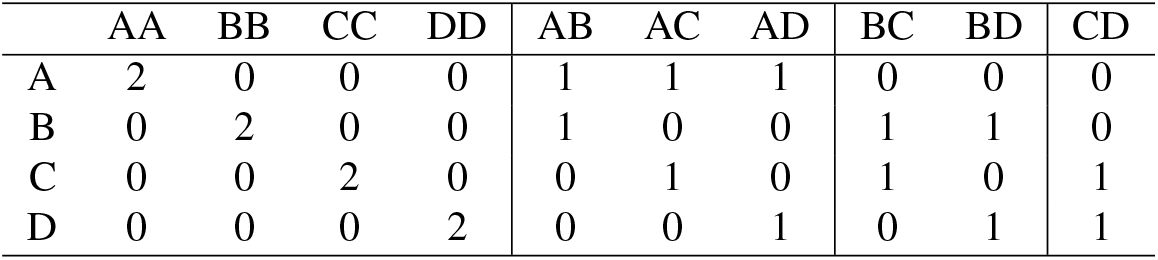
Summing matrix to transform allelic compositions into genotypic compositions.

|  | AA | BB | CC | DD | AB | AC | AD | BC | BD | CD |
| --- | --- | --- | --- | --- | --- | --- | --- | --- | --- | --- |
| A | 2 | 0 | 0 | 0 | 1 | 1 | 1 | 0 | 0 | 0 |
| B | 0 | 2 | 0 | 0 | 1 | 0 | 0 | 1 | 1 | 0 |
| C | 0 | 0 | 2 | 0 | 0 | 1 | 0 | 1 | 0 | 1 |
| D | 0 | 0 | 0 | 2 | 0 | 0 | 1 | 0 | 1 | 1 |

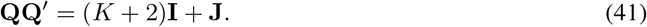

Let 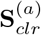and 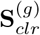be the covariance matrices of clr transformed allelic and genotypic compositions respec-tively. We have

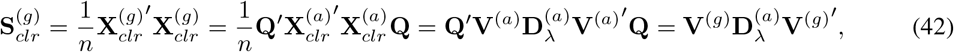

where 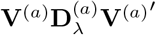 is the spectral decomposition of 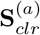. The total genotypic variance is given by

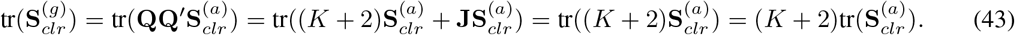

We have that

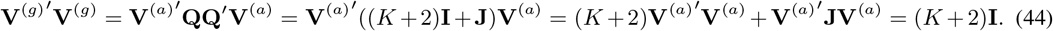

Scaling these eigenvectors to unit norm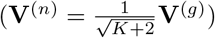, we have

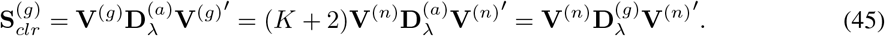

which shows

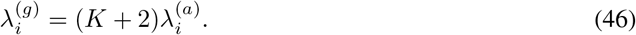

It follows that the eigenvectors obtained in the genotypic analysis satisfy

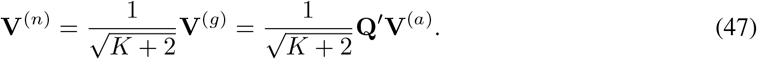

In terms of the SVD for the LR-PCA (see Eq. 13) of the genotypic data, we have

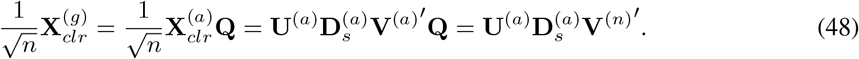

It follows that for the genotypic analysis the principal components are obtained as

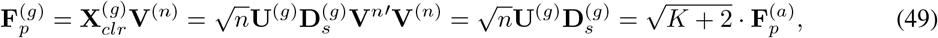

and biplot vectors, in standard coordinates, for the clr parts are given by

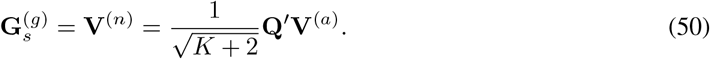

## Supplementary figures

**Figure S1:**
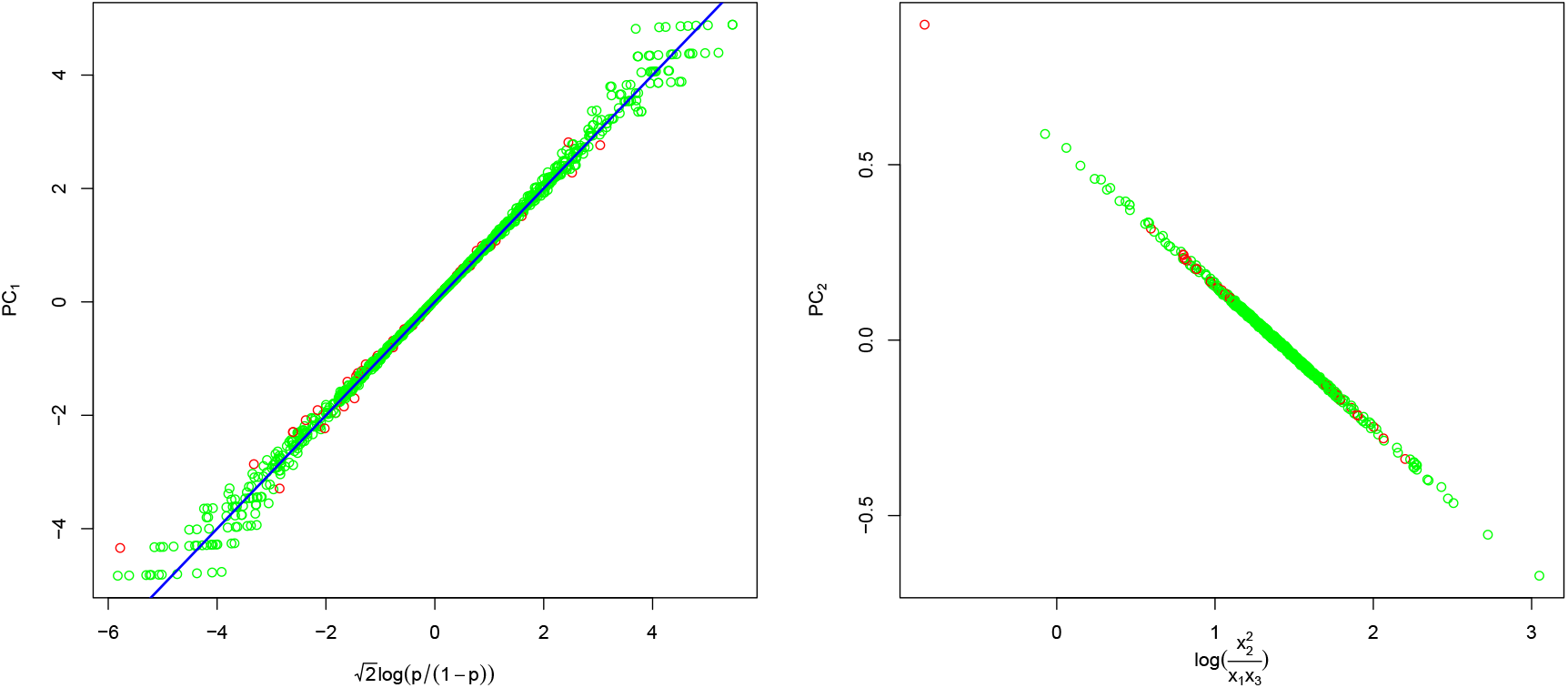
Interpretation of principal components for multiple samples from a biallelic polymorphism. A: Scatterplot of the first principal component against the scaled logit of the allele frequency. B: Scatterplot of the second principal component against the disequilibrium measure.

**Figure S2:**
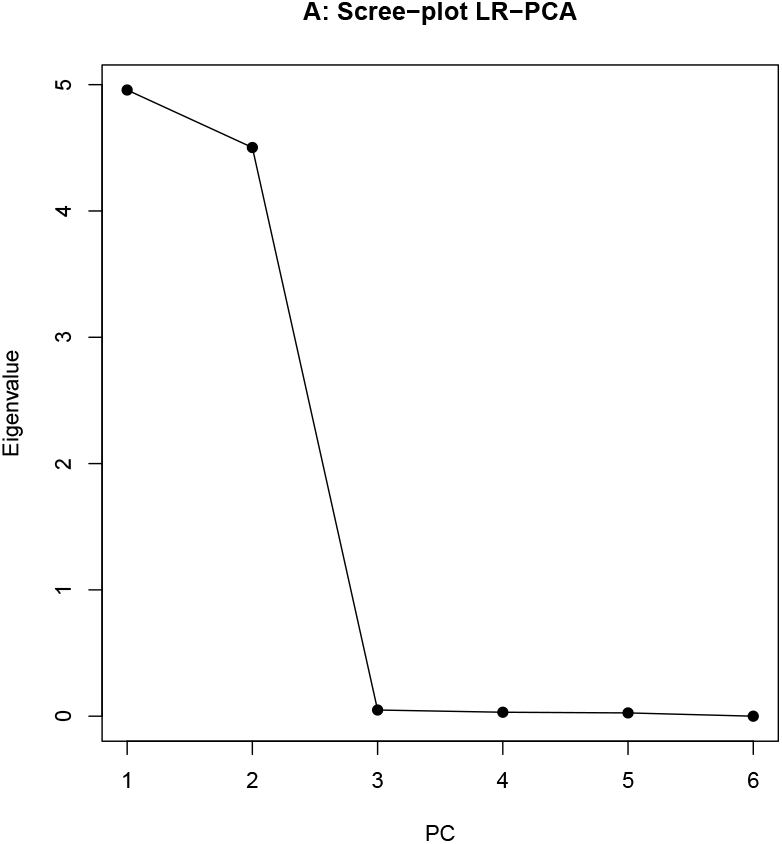
Screeplot of eigenvalue of LR-PCA with simulated samples for triallelic variant under HWE.

**Figure S3:**
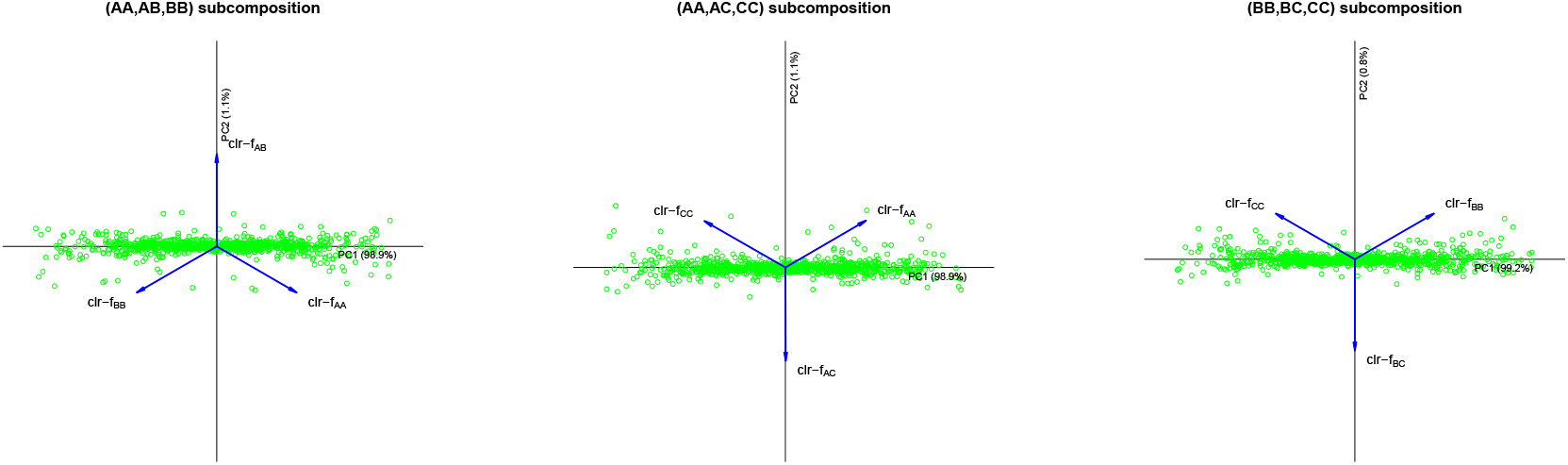
LR-PCA biplots with two-allelic subcompositions of a triallelic variant under HWE.

**Figure S4:**
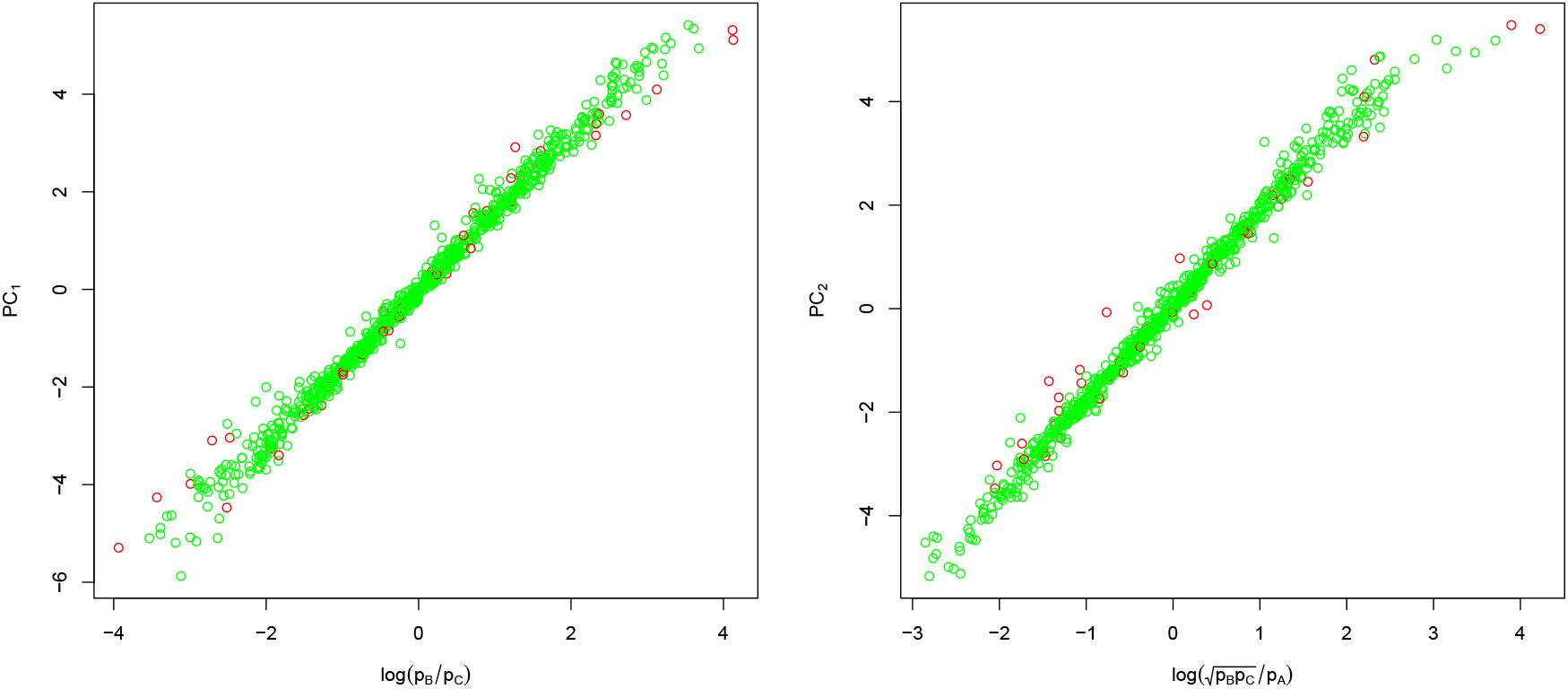
Scatterplots of the first two principal components against balances of allele frequencies for a triallelic variant under HWE.

**Figure S5:**
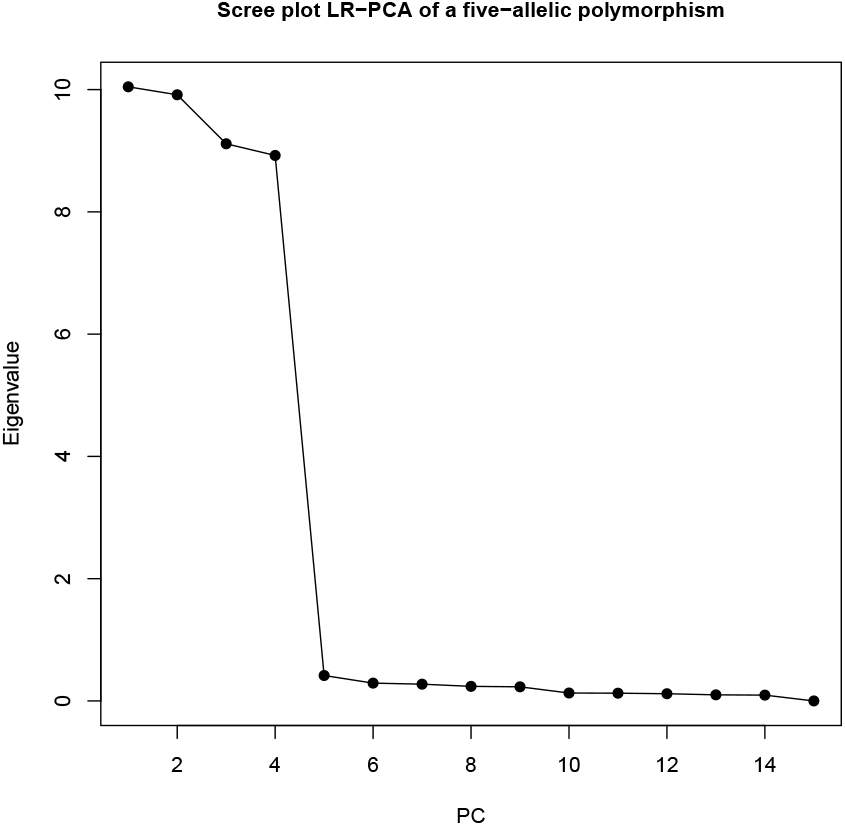
Scree plot of the LR-PCA of a five-allelic variant under HWE.

**Figure S6:**
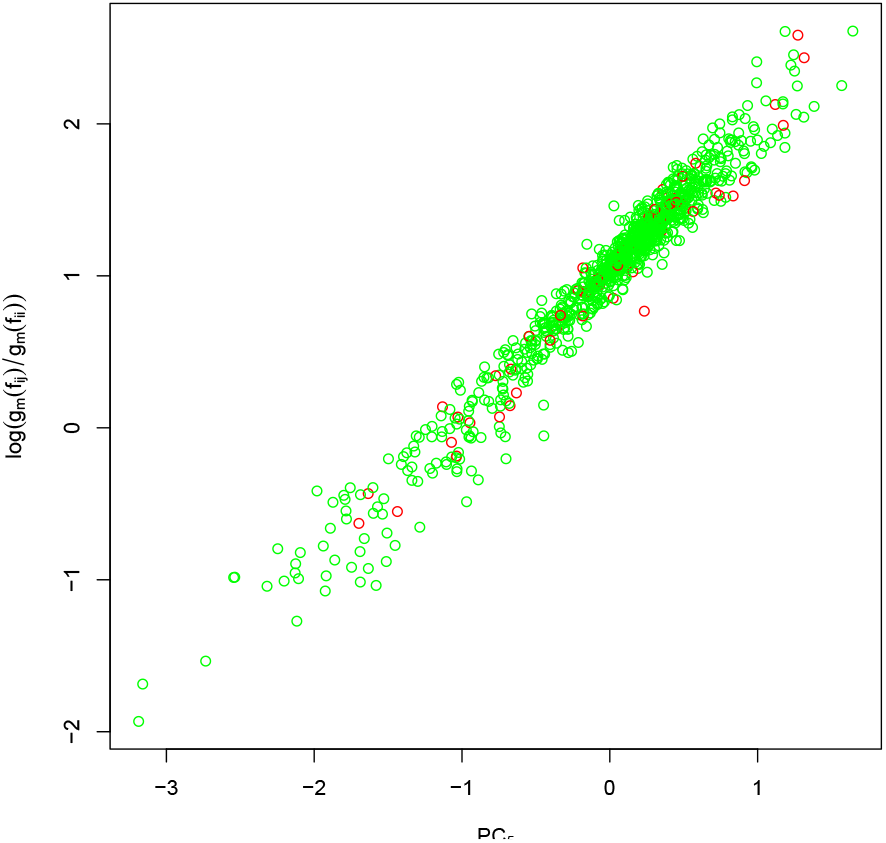
Scatterplot of *PC*_5_ obtained in the LR-PCA of a five-allelic variant under HWE against the heterozygote-homozygote balance.

**Figure S7:**
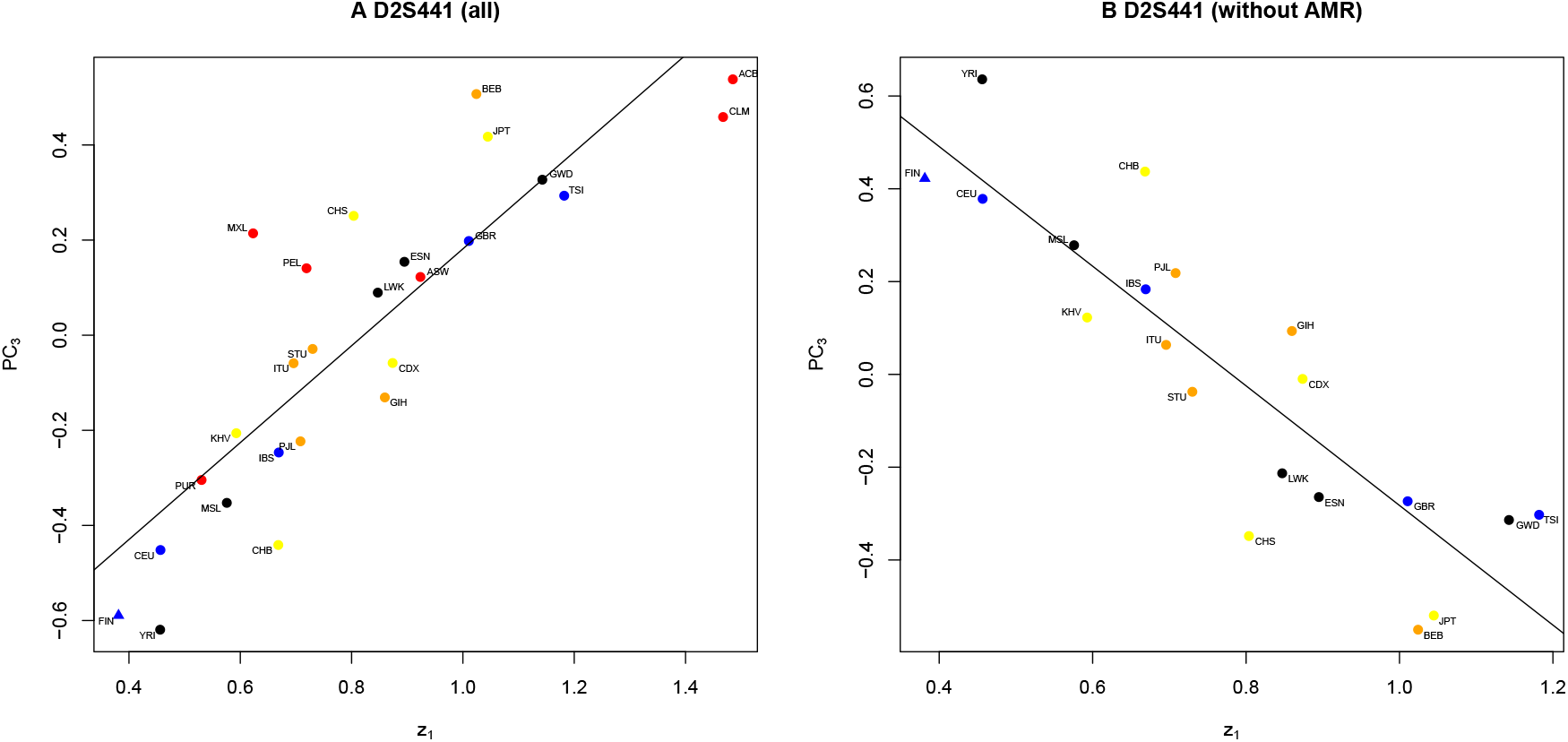
Scatterplots of *PC*_3_ against disequilibrium balance *z*_1_ obtained in the LR-PCA of triallelified STR D2S441.

**Table S1:**
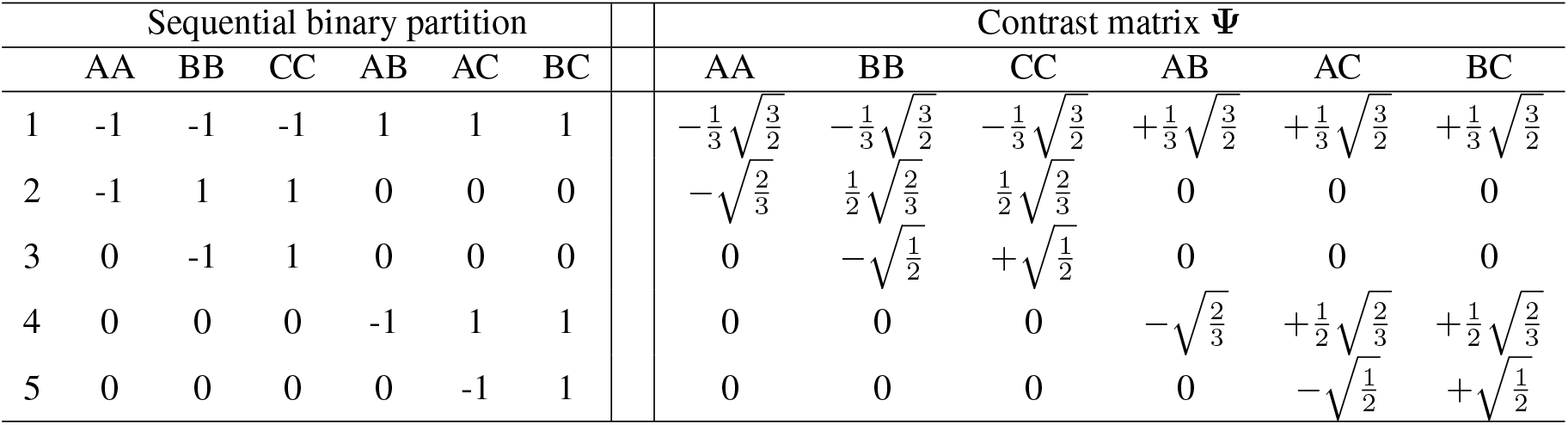
Sequential binary partition and contrast matrix **Ψ** for a triallelic autosomal polymorphism.

| Sequential binary partition | | | | | | | Contrast matrix $\Psi$ | | | | | |
| --- | --- | --- | --- | --- | --- | --- | --- | --- | --- | --- | --- | --- |
|  | AA | BB | CC | AB | AC | BC | AA | BB | CC | AB | AC | BC |
| 1 | -1 | -1 | -1 | 1 | 1 | 1 | $-\frac{1}{3}\sqrt{\frac{3}{2}}$ | $-\frac{1}{3}\sqrt{\frac{3}{2}}$ | $-\frac{1}{3}\sqrt{\frac{3}{2}}$ | $+\frac{1}{3}\sqrt{\frac{3}{2}}$ | $+\frac{1}{3}\sqrt{\frac{3}{2}}$ | $+\frac{1}{3}\sqrt{\frac{3}{2}}$ |
| 2 | -1 | 1 | 1 | 0 | 0 | 0 | $-\sqrt{\frac{2}{3}}$ | $\frac{1}{2}\sqrt{\frac{2}{3}}$ | $\frac{1}{2}\sqrt{\frac{2}{3}}$ | 0 | 0 | 0 |
| 3 | 0 | -1 | 1 | 0 | 0 | 0 | 0 | $-\sqrt{\frac{1}{2}}$ | $+\sqrt{\frac{1}{2}}$ | 0 | 0 | 0 |
| 4 | 0 | 0 | 0 | -1 | 1 | 1 | 0 | 0 | 0 | $-\sqrt{\frac{2}{3}}$ | $+\frac{1}{2}\sqrt{\frac{2}{3}}$ | $+\frac{1}{2}\sqrt{\frac{2}{3}}$ |
| 5 | 0 | 0 | 0 | 0 | -1 | 1 | 0 | 0 | 0 | 0 | $-\sqrt{\frac{1}{2}}$ | $+\sqrt{\frac{1}{2}}$ |

**Table S2:** Eigenvectors obtained in an ilr based LR-PCA for a triallelic autosomal polymorphism. The relation between the heterozygote-homozygote balance with the third component is shown in bold.

|  | PC1 | PC2 | PC3 | PC4 | PC5 |
| --- | --- | --- | --- | --- | --- |
| $z_1$ | 0.0018 | -0.0021 | **-0.9830** | 0.0362 | -0.1798 |
| $z_2$ | 0.0459 | 0.8867 | -0.0837 | 0.0201 | 0.4520 |
| $z_3$ | 0.8882 | -0.0430 | 0.0226 | 0.4564 | -0.0221 |
| $z_4$ | 0.4063 | 0.2062 | 0.0447 | -0.7929 | -0.4022 |
| $z_5$ | -0.2097 | 0.4116 | 0.1553 | 0.4016 | -0.7753 |

**Table S3:** Eigenvectors obtained in the clr based LR-PCA of a five-allelic autosomal polymorphism under HWE, after zero-imputation. Coefficients of the fifth eigenvector are emphasised.

|  | PC1 | PC2 | PC3 | PC4 | PC5 | PC6 | PC7 | PC8 | PC9 | PC10 | PC11 | PC12 | PC13 | PC14 |
| --- | --- | --- | --- | --- | --- | --- | --- | --- | --- | --- | --- | --- | --- | --- |
| AA | 0.4611 | 0.4111 | 0.1329 | -0.0549 | **0.3528** | 0.2147 | 0.1275 | 0.5695 | -0.1089 | 0.0197 | 0.0675 | 0.0358 | -0.0137 | 0.0412 |
| BB | 0.2832 | -0.5119 | -0.2251 | -0.1052 | **0.2883** | -0.4174 | 0.4789 | -0.0878 | -0.1556 | -0.0225 | -0.0757 | 0.0076 | 0.0120 | 0.0663 |
| CC | -0.1345 | -0.1178 | 0.3126 | 0.5217 | **0.2731** | -0.0520 | 0.0999 | -0.0002 | 0.6522 | 0.1018 | 0.0526 | 0.0018 | -0.0522 | -0.0542 |
| DD | -0.3287 | -0.0187 | 0.2772 | -0.4473 | **0.4823** | -0.2501 | -0.4798 | -0.0645 | -0.1065 | 0.0627 | 0.0266 | -0.0380 | 0.0322 | -0.0048 |
| EE | -0.2693 | 0.2439 | -0.5098 | 0.0957 | **0.3663** | 0.4573 | 0.1392 | -0.3992 | -0.0882 | 0.0310 | 0.0211 | -0.0106 | 0.0013 | 0.0298 |
| AB | 0.4307 | -0.0644 | -0.0522 | -0.0942 | **-0.1488** | 0.1282 | -0.2988 | -0.2369 | 0.1466 | 0.1328 | -0.1861 | -0.4915 | -0.4798 | -0.0508 |
| AC | 0.1959 | 0.1641 | 0.2520 | 0.2644 | **-0.1278** | -0.0979 | -0.0150 | -0.3140 | -0.3030 | -0.0485 | 0.0560 | -0.2318 | 0.5437 | -0.4040 |
| BC | 0.0897 | -0.3619 | 0.0515 | 0.2402 | **-0.0759** | 0.2813 | -0.3497 | 0.0486 | -0.1361 | -0.3727 | -0.0076 | 0.0926 | 0.2119 | 0.5599 |
| AD | 0.0715 | 0.2213 | 0.2430 | -0.3057 | **-0.2833** | -0.0485 | 0.2515 | -0.3530 | 0.1418 | -0.0992 | 0.5319 | 0.2079 | -0.1340 | 0.2870 |
| BD | -0.0393 | -0.3090 | 0.0400 | -0.3226 | **-0.2501** | 0.3916 | 0.0734 | 0.1132 | 0.1354 | 0.5477 | -0.1119 | 0.2043 | 0.3259 | -0.1336 |
| CD | -0.2766 | -0.0786 | 0.3475 | 0.0412 | **-0.1360** | 0.1789 | 0.2314 | 0.0362 | -0.3248 | -0.2856 | -0.3398 | 0.2069 | -0.4453 | -0.2815 |
| AE | 0.1065 | 0.3725 | -0.2195 | 0.0158 | **-0.1601** | -0.3494 | -0.1674 | -0.0797 | 0.2056 | -0.0276 | -0.5365 | 0.4529 | 0.0951 | 0.0877 |
| BE | 0.0004 | -0.1502 | -0.4131 | -0.0015 | **-0.1228** | -0.0668 | -0.2697 | 0.2513 | 0.1169 | -0.2832 | 0.4391 | 0.1912 | -0.0867 | -0.5039 |
| CE | -0.2326 | 0.0747 | -0.1129 | 0.3562 | **-0.2264** | -0.2787 | -0.0593 | 0.2048 | -0.3813 | 0.5236 | 0.1965 | -0.0637 | -0.2127 | 0.2282 |
| DE | -0.3578 | 0.1250 | -0.1240 | -0.2040 | **-0.2314** | -0.0912 | 0.2380 | 0.3118 | 0.2058 | -0.2801 | -0.1338 | -0.5654 | 0.2022 | 0.1328 |

## Notes

### Competing Interest Statement

The authors have declared no competing interest.

